# Quantification of esterified oxylipins following HILIC-fractionation of lipid classes

**DOI:** 10.1101/2025.09.09.675214

**Authors:** Luca M. Wende, Laura Carpanedo, Lilli Scholz, Nadja Kampschulte, Annette L. West, Philip C. Calder, Nils Helge Schebb

**Author notes:** corresponding author: Nils Helge Schebb, Food Chemistry, School of Mathematics and Natural Sciences, University of Wuppertal, Gausstrasse 20, 42119 Wuppertal, Germany.

## Abstract

Several oxylipins are lipid mediators derived from the oxidation of polyunsaturated fatty acids (PUFAs). The majority of oxylipins in biological samples occurs esterified in neutral lipids (nLs) and phospholipids (PLs). They are commonly quantified indirectly following alkaline hydrolysis providing excellent sensitivity but the information in which lipid classes the oxylipins occurred in is lost. The direct analysis of oxidized lipids is currently not sensitive enough to detect all esterified oxylipins. Here, a new hydrophilic interaction liquid chromatography (HILIC) based lipid class fractionation using solid-phase extraction (SPE) cartridges was developed separating lipids into nLs and 4 PL fractions using a single column. Esterified oxylipins in the fractions were quantified following alkaline hydrolysis to sensitively pinpoint in which lipid classes they are bound in plasma. The fractionation was extensively characterized for different lipid extracts demonstrating high separation efficiency and recovery using labeled standards and untargeted analysis of endogenous lipids. Esterified oxylipins in the fractions were quantitatively detected. Based on the results from two independent human plasma pools including SRM1950 it is shown that: hydroxy-linoleic acid- and hydroxy-α-linolenic acid-derived oxylipins are preferably bound to nLs whereas long chain hydroxy-PUFAs and PUFAs (i.e. ARA EPA and DHA) are predominantly esterified to phospholipid classes. Supplementation of n3-PUFAs for 12 months led to an increase in EPA- and -DHA-derived oxylipins in all lipid fractions with the highest increase of hydroxy-PUFAs in nLs. This demonstrates a precursor PUFA-dependent binding of oxylipins and a direct effect of diet on esterified oxylipins in plasma.

## Introduction

Several eicosanoids and other oxylipins are potent lipid mediators derived from the oxidation of PUFAs (1). They can be formed either nonenzymatically by radical-mediated and stereo-random (aut)oxidation or enzymatically by different enzymes, leading to position- and stereo-specific products (2). The enzymatic pathways of formation play important roles in the regulation of inflammation, immunity, thrombosis and other biological responses (3). PUFAs such as ARA can be oxidized by enzymes including cyclooxygenases (COX), lipoxygenases (LOX), and cytochrome P450 monooxygenases (CYP) (4). LOX enzymes (e.g., 5-LOX, 12-LOX, 15-LOX-1, 15-LOX-2) form regio- and stereoselective hydroperoxy-PUFAs at different positions, which can be reduced to hydroxy-PUFAs by enzymes such as glutathione peroxidases (5, 6). The oxidation of PUFAs by CYP results in the formation of terminal hydroxy-PUFAs and/or cis-(*R*, *S* and *S*, *R*)-epoxy-PUFAs (7). Oxylipins can be formed from n6-PUFAs such as ARA and n3-PUFAs including EPA and DHA, giving rise to a large number of structurally diverse oxidized fatty acids (8).

The vast majority of oxylipins, i.e. epoxy-PUFAs and hydroxy-PUFAs present in blood plasma (9), serum (10), cells (11), and tissues (12) occur in esterified form. Similar to their PUFA precursors, oxylipins are bound as esters in lipids such as cholesterol esters (Chol Ester), triacylglycerols (TGs), and polar lipids, i.e., glycerophospholipids (PLs) (13). Indeed, >90% of the hydroxy-PUFAs present in plasma are esterified (14). Little is known about which lipids the oxylipins are esterified in but some studies suggest that they are predominantly bound to TGs and PLs (15). Notably, oxylipins such as 18:2;OH, and 20:4;OH were found esterified in PC, TG, and Chol Ester species in human plasma (16), as well as in PC species in human serum (17).

Analysis of esterified oxylipins is challenging due to their low abundances in biological samples. Esterified oxylipins can be analyzed directly as oxidized PLs or TGs by LC-MS, providing detailed information about the binding form and lipid species of the bound oxylipin (13, 18). Quantitative targeted LC-MS/MS of oxPLs (bearing oxylipins) showed a distinct incorporation of hydroxy-PUFAs in both, oxidized PL classes and species, in human cells, but such research has been limited to a few oxylipins bound in selected oxidized PLs (11, 19, 20)

Even targeted methods (19, 20) using the most sensitive instrumentation available are limited by the sensitivity because the low concentrations of oxylipins are distributed over a large number of lipid classes and species including neutral lipids (nLs) such as TG, Chol Ester and PL classes such as phosphatidyl-choline (PC), -ethanolamine (PE), -glycerol (PG), -inositol (PI), and -serine (PS). This is highlighted in (Fig. 1) demonstrating as an example for 15-HETE the vast number of possible binding forms. Moreover, the direct analysis of esterified oxylipins is limited by the lack of available standards hindering identification and quantification (11, 16, 21). Thus, quantification of esterified oxylipins in biological samples is commonly performed indirectly by alkaline hydrolysis followed by quantification of the resulting free oxylipins using established targeted LC-MS/MS methods (10, 14, 22–24). However, the information about which lipid class the oxylipins are bound in gets lost in the saponification step. To better understand in which lipid classes esterified oxylipins are bound, we developed a sensitive orthogonal approach: lipid class fractionation using a solid-phase extraction (SPE) cartridge followed by LC-MS/MS quantification of esterified oxylipins in the different lipid fractions after alkaline hydrolysis. Chromatographic fractionation of lipids using SPE cartridges is frequently applied, allowing the separation of Chol Ester, TGs, and different PLs such as PC, PE, PG, PI/PS classes (25–27). However, for oxylipins, this was only used so far to differentiate between nLs and PLs in cells (28) and plasma (29). Here, we utilize a not previously reported hydrophilic interaction liquid chromatography (HILIC) approach for separation requiring only a single SPE column to efficiently fractionate lipids in nLs and four different PL fractions. The performance of the fractionation was characterized for different biological samples using non-targeted lipidomics. The method was then applied to the investigation of esterified oxylipins in human plasma and the changes induced by n3-PUFA supplementation. A distinct PUFA-dependent binding of hydroxy-PUFAs in plasma lipids was unveiled.

**Fig. 1.**
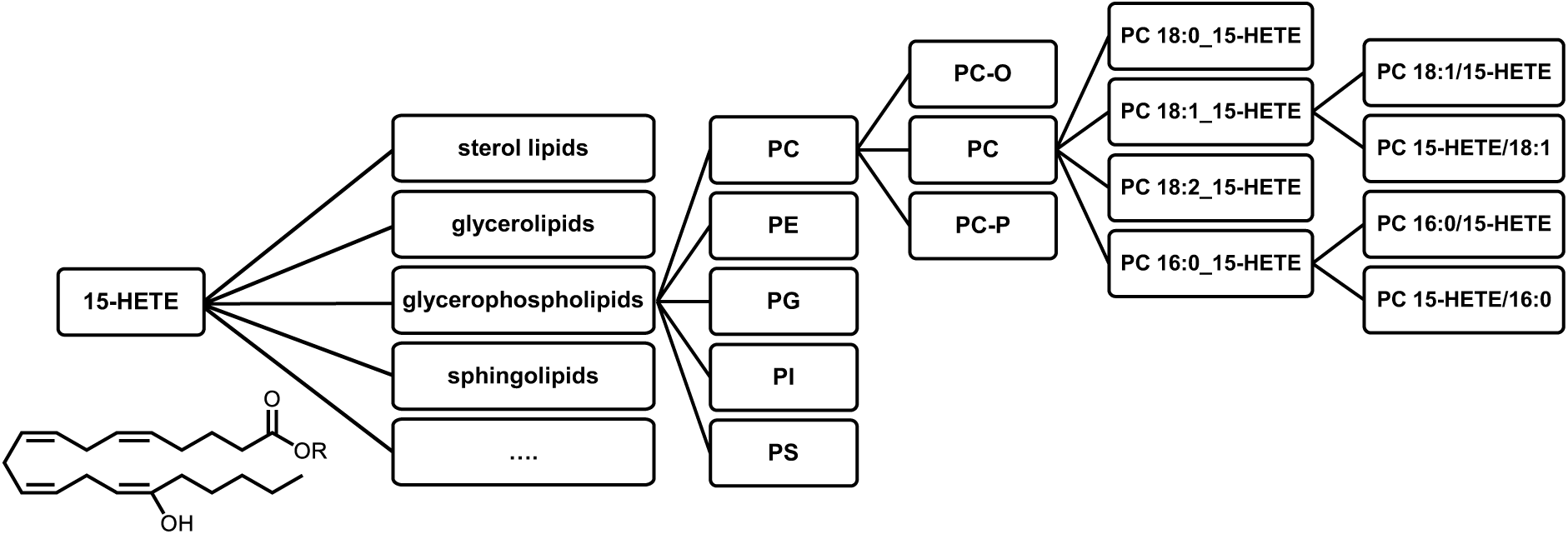
Distribution of esterified oxylipin across different lipid classes and species. This illustration highlights for 15-HETE and PLs the large number of different binding forms of esterified oxylipins.

## Materials and methods

### Chemicals and biological material

The internal standard mixtures EquiSPLASH Lipidomix (330731-1EA) containing PC 15:0/18:1[D7], lyso-PC 18:1[D7]/0:0, PE 15:0/18:1[D7], lyso-PE 18:1[D7]/0:0, PG 15:0/18:1[D7], PI 15:0/18:1[D7], PS 15:0/18:1[D7], TG 15:0/18:1[D7]/15:0, diacylglycerol (DG) 15:0/18:1[D7]/0:0, monoacylglycerol (MG) 18:1[D7]/0:0/0:0, Chol Ester 18:1[D7], SM 18:1;O2/18:1[D9], C15 Ceramide (Cer)[D7] (each 100 µg/mL), and SPLASH Lipidomix (30707-1EA) containing PC 15:0/18:1[D7], lyso-PC 18:1[D7]/0:0, PE 15:0/18:1[D7], lyso-PE 18:1[D7]/0:0, PG 15:0/18:1[D7], PI 15:0/18:1[D7], PS 15:0/18:1[D7], TG 15:0/18:1[D7]/15:0, DG 15:0/18:1[D7]/0:0, MG 18:1[D7]/0:0/0:0, Chol Ester 18:1[D7], sphingomyelin (SM) 18:1;O2/18:1[D9], PA 15:0/18:1[D7], and free cholesterol[D7] (concentrations 1.8 µg/mL ‒ 329.1 µg/mL) were purchased from Avanti Polar Lipids (local supplier: Merck KGaA, Darmstadt, Germany).

Acetonitrile (ACN) LC-MS grade, methanol (MeOH) LC-MS grade, isopropanol (IPA) LC-MS grade, formic acid LC-MS grade, and acetic acid LC-MS grade were obtained from Fisher Scientific (Schwerte, Germany). Ultra-pure water was generated using a Barnstead Genpure Pro system from Thermo Fisher Scientific (Langenselbold, Germany).

Human embryonic kidney 293 (HEK293) cells were obtained from the German Collection of Microorganisms and Cell Cultures GmbH (DSMZ, Braunschweig, Germany).

Citrated blood plasma samples (plasma pool 1) were generated from three healthy adults approved by the ethics committee of the University of Wuppertal. Written informed consent was obtained from all subjects in accordance with the Declaration of Helsinki. In addition, human plasma reference material (National Institute of Standards and Technology (NIST) SRM 1950: Metabolites in Frozen Human Plasma) was analyzed.

### Cell culture

HEK293 cells were cultured in DMEM with high glucose (4.5 g/L), supplemented with 10% (v/v) FCS, 100 U/mL penicillin, 100 μg/mL streptomycin, and 1 mM sodium pyruvate. Culture occurred in a humidified atmosphere containing 5% CO_2_ at 37 °C. Cells were seeded at a density of 5 × 10^6^ cells per 60.1 cm^2^ dish and harvested by scraping in ice-cold PBS after 24 h.

### Sample preparation

As illustrated in Fig. 2, 100 µL human plasma or 100 µL sonicated cell suspension (⁓400 µg cell protein) in H_2_O/MeOH (50:50, v/v) were transferred into a 1.5 mL microcentrifuge tube, followed by the addition of 10 µL of an internal standard (IS) mixture (EquiSPLASH for cell suspensions [2 µL/⁓400 µg cell protein], SPLASH Lipidomix for human plasma [5 µL/100 µL plasma], or deuterium-labeled oxylipins [each 1 pmol/100 µL plasma/⁓400 µg cell protein]). 10 µL of inhibitor/antioxidant solution (0.2 mg/mL butylated hydroxytoluene, 100 mM indomethacin, and 100 mM of the soluble epoxide hydrolase inhibitor trans-4-[4-(3-adamantan-1-yl-ureido)-cyclohexyloxy]-benzoic acid) was added. Protein precipitation was performed using 400 µL of ice-cold IPA and samples were mixed and frozen at – 80°C for at least 30 min. After centrifugation (20,000 × g, 10 min, 4 °C), the supernatant was used for SPE-based lipid class fractionation or quantification of oxylipins/FA without prior lipid fractionation.

**Fig. 2.**
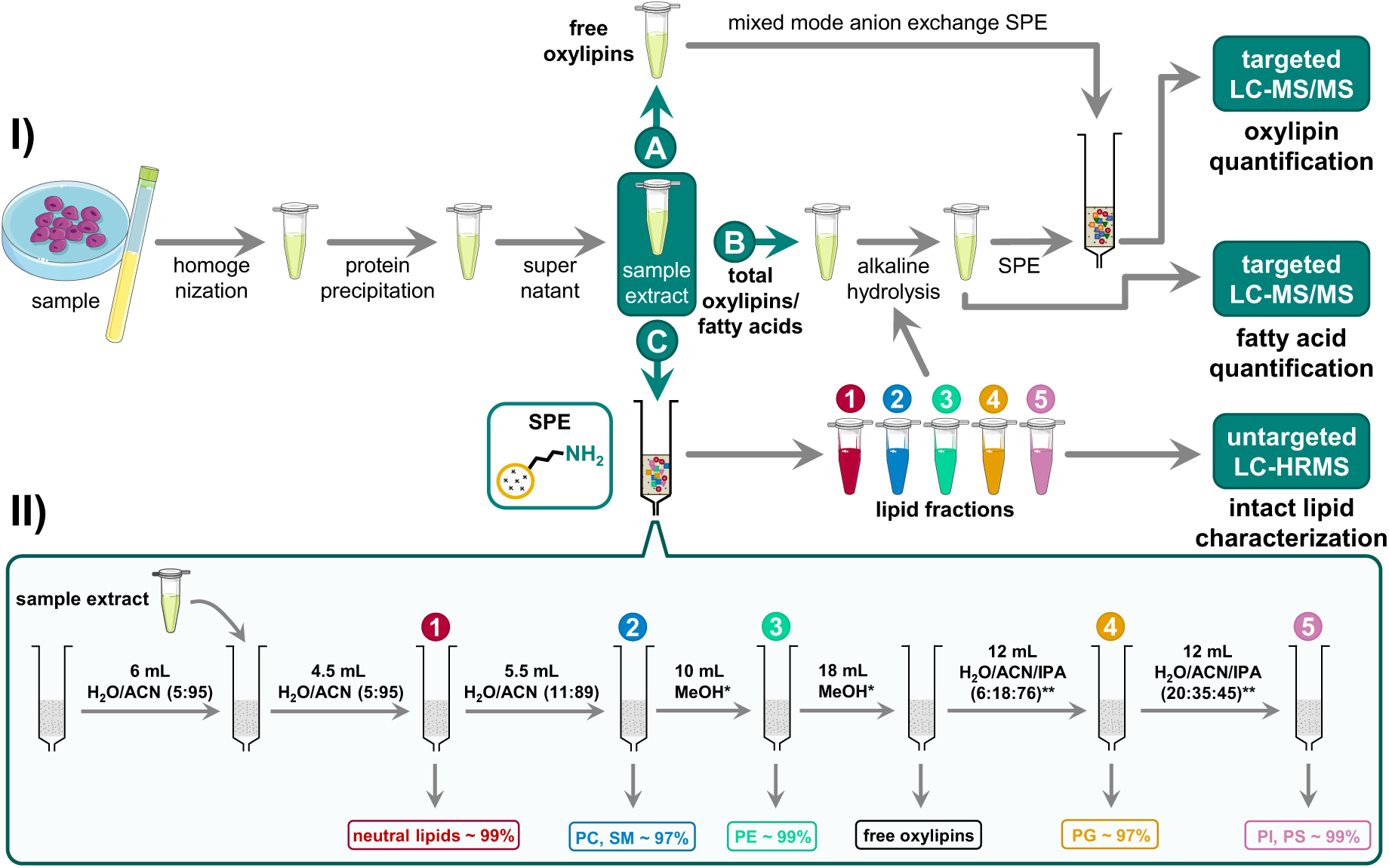
Scheme of the lipid class-specific analysis of esterified FA and oxylipins. **I)** Samples are homogenized, and proteins are precipitated. The supernatant is used for **(A)** the direct analysis of non-esterified oxylipins, **(B)** total oxylipins/FA after alkaline hydrolysis, and **(C)** total FA and oxylipin analysis following fractionation of lipid classes and alkaline hydrolysis. **II)** Lipid classes are fractionated by HILIC-based SPE into five fractions using a single column. **II)** The recovery of the lipids (determined for SPLASH ISs) in the fractions is indicated. *0.1% acetic acid; **20mM ammonium formate + 0.1% formic acid.

### Fractionation of lipid classes using a solid-phase extraction cartridge

SPE for lipid class fractionation was performed using CHROMABOND NH_2_ cartridges (3 mL/500 mg sorbent per cartridge, particle size 45 µm; Machery-Nagel, Düren, Germany). Cartridges were washed with two volumes of MeOH and two volumes of ACN and preconditioned with 6 mL of H_2_O/ACN (5:95, v/v) (Fig. 2C). Then 1000 µL of ACN, together with 500 µl of the sample after protein precipitation, were added onto the column. nLs including MGs, DGs, TGs, Chol Ester, and Cers were eluted with 4.5 mL of H_2_O/ACN (5:95, v/v). The second fraction, containing all choline-bearing PLs (i.e., lyso-PCs, PCs, and SMs), was collected with 5.5 mL of H_2_O/ACN (11:89, v/v). The third fraction, comprising lyso-PEs and PEs, was isolated using 10 mL of MeOH with 0.1% acetic acid. Subsequently, 18 mL of MeOH with 0.1% acetic acid was used to remove free oxylipins and FAs. PGs were eluted with 12 mL of H_2_O/ACN/IPA (6:18:76, v/v/v), and PIs and PSs were simultaneously isolated using 12 mL of H_2_O/ACN/IPA (20:35:45, v/v/v). Both solvents for PG and PI/PS elution contained 20 mM ammonium formate and 0.1% formic acid. Individual lipid fractions were eluted under a slight vacuum (∼900 mbar) and collected in borosilicate glass tubes containing 8 µL of 30% glycerol in MeOH. After collection, each fraction was spiked with deuterium-labeled oxylipin internal standards (each 1 pmol/100 µL plasma/cell extract (∼400 µg cell protein)) and solvents were evaporated using a CHRIST CT 02-50 rotary evaporator (30° C, 1 mbar). The residues were reconstituted in 100 µL ACN/IPA (50:50, v/v). Lipid extracts were either analyzed by non-targeted LC-high-resolution-MS (LC-HRMS) (30) for method characterization described in the supplementary information or prepared for the quantification of total oxylipins/FAs. For this, the reconstituted fractions were mixed with 300 µL IPA and centrifuged. Hydrolysis was carried out following the addition of 100 µl water and 100 µL 0.6 M KOH in H_2_O/MeOH 25:75, v/v) for 30 min at 60 °C.

### Quantification of oxylipins and fatty acids

The analysis of free and total oxylipins was conducted as described elsewhere (23, 31–34). Briefly, after protein precipitation in organic solvent, oxylipins were extracted either directly (free oxylipins, Fig. 2A), or after alkaline hydrolysis (23) (total oxylipins, Fig. 2B) using mixed-mode anion exchange Oasis Max cartridges, (3 mL, 60 mg sorbent per cartridge, particle size 30 µm, from Waters, Eschborn, Germany. LC-MS/MS oxylipin analysis was performed by external calibration with internal standards using a 1290 Infinity LC II System (Agilent Technologies, Waldbronn, Germany) coupled to a QTRAP5500 mass spectrometer operating in scheduled selected reaction monitoring mode. Fatty acids were quantified using the same sample preparation as described elsewhere (35). LC-MS/MS fatty acid analysis was performed by external calibration with internal standards using a 1260 Infinity LC System (Agilent Technologies, Waldbronn, Germany) coupled to an API 3200 instrument (AB Sciex, Darmstadt, Germany) operating in scheduled selected reaction monitoring mode.

### n3-PUFA supplementation study

The effect of n3-PUFA supplementation on the esterification of oxylipins in different lipid classes was investigated using human plasma samples derived from a double-blinded, randomized, controlled intervention trial (36). A subset of 9 subjects (3 males, 6 females, age 23 ‒ 72 years) was selected (Supplemental Fig. S1). Four times a week, each participant received n3-PUFA capsules containing EPA and DHA (1.5 g and 1.8 g, respectively) from fish oil as re-esterified TG equivalent to 4 portions of fatty fish per week. Plasma samples of the subjects at baseline level and after 12 months of n3-PUFA supplementation were analyzed.

## Results

Lipids were fractionated into nLs and four different PL classes (i.e., PC, PE, PG, and PI/PS), enabling the analysis of the binding form of oxylipins and their precursor PUFAs (Fig. 2). For this, a new SPE fractionation protocol using only a single cartridge was developed. The separation efficiency of each lipid class was evaluated using deuterium-labeled lipids (SPLASH) spiked to plasma and cell samples before lipid fractionation (Table 1). Overall, high separation efficiency and minimal cross-contamination (<5%) were achieved for most spiked deuterium-labeled lipid standards. In fraction 1 nLs were isolated, including TG 15:0/18:1[D7]/15:0 (100%), DG 15:0/18:1[D7]/0:0 (100%), MG 18:1[D7]/0:0/0:0 (100%), and Chol Ester 18:1[D7] (91%); no contamination with PLs was observed. Fraction 2 contained choline-bearing PLs, such as PC 15:0/18:1[D7] (100%), lyso-PC 18:1[D7]/0:0 (60%), and SM 18:1;O2/18:1[D9] (97%). Fraction 3 comprised PE 15:0/18:1[D7] (100%), lyso-PE 18:1[D7]/0:0 (100%), and 40% of lyso-PC 18:1[D7]/0:0. Free oxylipins and free FAs were quantitatively eluted from the cartridge with 18 mL MeOH + 0.1% acetic acid (Supplemental Fig. S2). In fraction 4, PG 15:0/18:1[D7] (95%) was isolated. In fraction 5 both PS 15:0/18:1[D7] (100%) and PI 15:0/18:1[D7] (100%) were eluted. Of note, the elution of PG, PI, and PS required a minimum concentration of 10 mM ammonium formate (Supplemental Fig. S3).

**Table 1.** Separation efficiency of the lipid class fractionation by SPE. A mixture of isotopically labeled lipid standards (SPLASH) was spiked to plasma and separated into five fractions. Shown are the relative recoveries of each standard in each fraction, reflecting the efficiency of lipid class separation across the fractions. Analysis was carried out using untargeted RP-LC-ESI-HRMS in Full MS/ddMS^2^ TOP N mode. Data is shown as mean ± SD (n = 3).

| Lipid Species | Fraction 1 | Fraction 2 | Fraction 3 | Fraction 4 | Fraction 5 |
| --- | --- | --- | --- | --- | --- |
| Chol Ester 18:1[D7] | 91 ± 2 | 4 ± 1 | 2.7 ± 0.4 | 1.6 ± 0.2 | 0.2 ± 0.1 |
| TG 15:0/18:1[D7]/15:0 | 99.6 ± 0.2 | 0.2 ± 0.1 | 0.2 ± 0.1 | 0.1 ± 0.1 | 0.1 ± 0.1 |
| DG 15:0/18:1[D7]/0:0 | 99.9 ± 0.1 | 0.1 ± 0.1 | n.d. | n.d. | n.d. |
| MG 18:1[D7]/0:0/0:0 | 99.9 ± 0.1 | n.d. | n.d. | n.d. | n.d. |
| PC 15:0/18:1[D7] | n.d. | 99.8 ± 0.2 | 0.2 ± 0.2 | n.d. | n.d. |
| SM 18:1;O2/18:1[D9] | n.d. | 97 ± 2 | 2.7 ± 1.6 | n.d. | n.d. |
| Lyso-PC 18:1[D7]/0:0 | n.d. | 45 ± 5 | 55 ± 5 | n.d. | n.d. |
| PE 15:0/18:1[D7] | n.d. | 0.1 ± 0.1 | 99.9 ± 0.1 | 0.1 ± 0.1 | n.d. |
| Lyso-PE 18:1[D7]/0:0 | n.d. | n.d. | 99.6 ± 0.1 | 0.4 ± 0.1 | 0.1 ± 0.1 |
| PG 15:0/18:1[D7] | n.d. | 0.3 ± 0.1 | 4 ± 1 | 95 ± 1 | 1.3 ± 0.1 |
| PI 15:0/18:1[D7] | n.d. | n.d. | n.d. | n.d. | 99.9 ± 0.1 |
| PS 15:0/18:1[D7] | n.d. | n.d. | n.d. | n.d. | 99.9 ± 0.1 |

The robustness of the separation process was shown on three different days and for different biological samples, i.e. human plasma and HEK293 cell extract (Supplemental Table S1). None of the tested matrices had a negative influence on the separation efficiency. When comparing the amount of recovered lipid standards added before fractionation (pre-spiked) and after the fractionation (post-spiked), more than 80% of each IS was recovered (Supplemental Fig. S4).

To demonstrate that the developed SPE protocol also enables the separation of endogenous lipids in human plasma and HEK293 cells, MS-DIAL was used for data analysis following untargeted LC-HRMS analysis of intact lipids. Based on the detected features (Supplemental Fig. S5 and Tables S2-S5), cumulative extracted ion chromatograms (XIC) were generated for each lipid class (TG, Chol Ester; PC, SM; lyso-PE, PE; PG; PS, PI). These were compared between the non-fractionated lipid extract (Fig. 3A) and fractionated lipid extracts of plasma and HEK293 cells (Fig. 3B, Supplemental Fig. S6). The cumulative XICs (peaks, retention time, and abundance) were similar between non-fractionated samples and the respective lipid class fraction (1–5) (Fig. 3A, B). This indicates that i) the lipids characterized in the non-fractionated samples are also present in the lipid class fraction and ii) endogenous lipids are efficiently separated into distinct fractions. Indeed, the peak pattern and the intensity of the XICs of fraction 1 (nLs) were identical to the non-fractionated plasma, supporting that all nLs were eluted in this fraction. Similarly, the chromatographic pattern of PC and SM species in the non-fractionated extract was recovered in fraction 2 (PC, SM), indicating a high recovery of the different choline-bearing PLs. In fraction 3 (PE), the cumulative XICs of PE features displayed a similar chromatographic pattern but with higher intensities compared to the non-fractionated sample. This is also reflected in a higher recovery of the IS PE 15:0/18:1[D7] (Fig. 3C): 110% in fraction 3 versus 65% in the non-fractionated sample. For the earlier eluting IS lyso-PE 18:1[D7]/0:0, a comparable recovery was found between fractionated and non-fractionated plasma extracts. This suggests that compounds co-eluting with PE – presumably abundant PC and SM species – suppressed the PE signals in the non-fractionated sample. A similar and even more striking matrix effect was observed for PG. While PG species were not detected in the non-fractionated human plasma lipid extract (Fig. 3A), several PG species with a characteristic peak pattern were detected in fraction 4 (Fig. 3B). The lack of detection in the non-fractionated plasma extract was partially caused by a co-elution of almost isobaric (< 3 ppm) M+1 ions of PE-O species (Supplemental Fig. S7). Again, the matrix effects were observed in the recovery of the IS PG 15:0/18:1[D7]: strong “ion enhancement” in the non-fractionated extract (recovery 370%) versus no effects in fraction 4 (recovery 110%) (Fig. 3C, Supplemental Fig. S8). Such “ion enhancement” effect of human plasma matrix on IS PG 15:0/18:1[D7] in RP-LC lipidomics has already been described (30). Fractionation also increased the abundance of PI/PS features in fraction 5 compared to the non-fractionated lipid extract. The recovery of the ISs PS 15:0/18:1[D7] and PI 15:0/18:1[D7] was also improved, indicating that PI/PS can be more sensitively detected following fractionation.

**Fig. 3.**
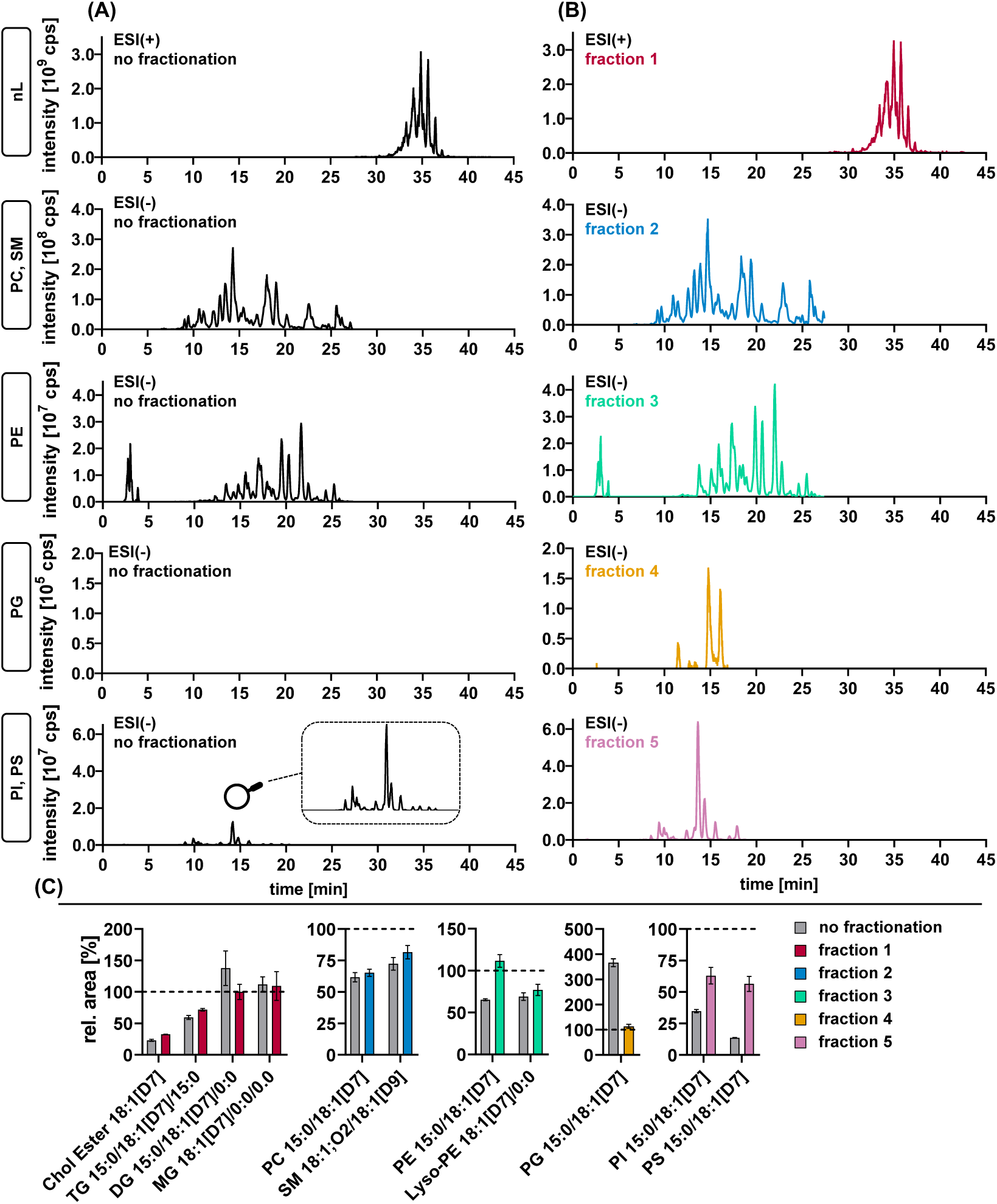
LC-HRMS lipidomics analysis of plasma with and without SPE-fractionation. Shown are cumulated XICs (sum of all detected species) of nLs and different PL classes in human plasma **(A)** without or **(B)** with fractionation. XICs were defined based on validated lipid features (lipid class level) using MS-DIAL (Supplemental Table S3). **(C)** Comparison of the recovery of ISs in the fractions to the non-fractionated sample showed a reduction of ion suppression effects for PE, PG, and PI/PS. ISs were spiked in plasma samples before extraction. Analysis was carried out using untargeted RP-LC-ESI-HRMS in Full MS/ddMS^2^ TOP N mode.

The SPE fractionation protocol was developed to quantitatively determine the distribution of oxylipins and PUFAs into lipid classes. For this purpose, oxylipins and FAs were released from each lipid fraction by an established alkaline hydrolysis procedure (22, 23, 33), and the concentrations of a comprehensive set of hydroxy-PUFAs were quantified in two pools of human plasma (Supplemental Tables S8 and S9). Figure 4 shows the concentrations of a representative set of hydroxy-PUFAs derived from LA, ALA, ARA, EPA, and DHA, along with their precursor PUFAs. The concentrations of total and free FAs and oxylipins were comparable to previous reports from human plasma (10, 24, 37, 38). LA-derived hydroxy-PUFA (HODEs) showed a manyfold higher concentration than all other oxylipins. In agreement with previous studies (14, 22, 39–41), the vast majority (≥85%) of hydroxy-PUFAs was detected in esterified form. The sum of FAs and oxylipins recovered from the SPE fractions closely matched the concentrations obtained in the non-fractionated human plasma samples. This indicates a high recovery of esterified FA/oxylipins and low loss or artificial formation/degradation during the fractionation procedure, enabling the evaluation of the distribution of the esterified oxylipins in the different lipid classes: Hydroxy-LAs and -ALAs (HOTrEs), and their precursors were predominantly esterified to nLs (60 – 86%). Hydroxy-ARAs (HETEs) and ARA were more prominently bound to PLs (66 – 70%). EPA, hydroxy-EPAs (HEPEs) and DHA were approximately equally distributed between nLs and PLs while Hydroxy-DHAs (HDHAs) occurred predominantly in PLs (75 – 86%).

**Fig. 4.**
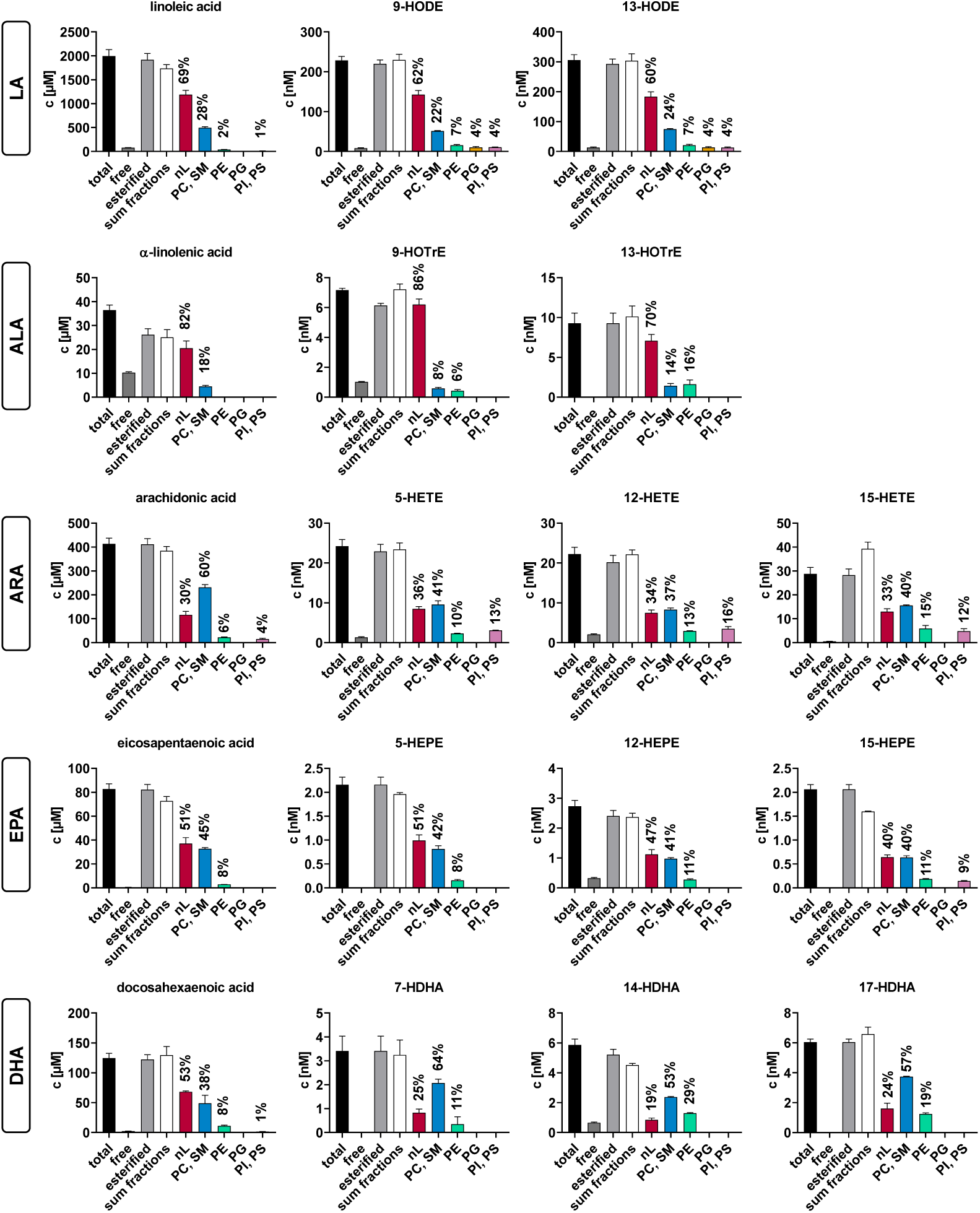
Lipid class-specific concentrations of esterified PUFAs and oxylipins in human plasma. Total FA and oxylipins in the five lipid fractions of human plasma (pool of 3 healthy subjects) were quantified by targeted RP-LC-ESI(-)-MS/MS following alkaline hydrolysis (Fig. 2C). A representative set of PUFAs and hydroxy-PUFAs from LA, ALA, ARA, EPA and DHA is shown in each fraction and compared to the results without fractionation (total and free FAs/oxylipins, Fig. 2A, B). The concentrations of all detected oxylipins can be found in Table S8. Shown are the concentrations ± SD (n = 3).

Among PL classes, LA, ALA, and their oxylipins were predominantly found in fraction 2, which included both PC and SM. Because SM species are composed of unsaturated or monounsaturated fatty acyl chains (42, 43), we conclude that these (hydroxy-)PUFAs are primarily bound to PC. In contrast, minor amounts were bound in PE, and only negligible amounts were found in PG and PI/PS. HETEs were predominantly esterified into PC (37 – 41%) followed by PE (10 – 15%), and PI/PS (12 – 16%). HEPEs showed comparable concentrations in nLs (40 – 51%) and in PC (40 – 44%) and were less esterified to PE (8 – 11%). Notably, 15-HEPE showed a relative concentration of 9% in PI/PS. The relatively high esterification into PI/PS was unique to HETEs and 15-HEPE. HDHAs, such as 7-and 14-HDHA were dominantly found in PC (53 – 64%) and PE (11 – 29%) but not detected in PG and PI/PS. Almost the same distribution of oxylipins was found in plasma pool 2 (NIST plasma, SRM 1950) (Supplemental Table S9).

Next, we investigated in which lipid classes the well-described changes (14, 44, 45) in esterified oxylipins occur following 12 months of n3-PUFA supplementation (Fig. 5). EPA and DHA supplementation did not change the concentration or distribution patterns of HODEs, HOTrEs, HETEs, and their n6-PUFA precursors (Supplemental Table S10). In contrast, n3-PUFA supplementation led to a marked increase in several HEPEs and HDHAs within the different lipid classes and shifted their distribution into lipids toward nLs. 5-HEPE and 12-HEPE increased from 2.5 ± 0.3 nM to 12 ± 2 nM, 15-HEPE from 2.2 ± 0.2 nM to 10 ± 1 nM, and 11-HEPE from 1.2 ± 0.2 nM to 3.8 ± 0.6 nM. The proportion of 5-HEPE in nLs was elevated from 53% to 62%, of 12-HEPE from 49% to 61%, of 15-HEPE from 51% to 61%, and of 11-HEPE from 55% to 63% after n3-PUFA supplementation. Following supplementation, 7-HDHA, 14-HDHA, 17-HDHA, and 13-HDHA showed concentrations in the same range as HEPEs (Fig. 5), despite higher baseline levels. 7-HDHA increased from 3.1 ± 0.3 nM to 10 ± 1 nM, 14-HDHA from 3.8 ± 0.3 nM to 7.8 ± 0.6 nM, 17-HDHA from 3.6 ± 0.3 nM to 8.9 ± 0.6 nM, and 13-HDHA from 3.6 ± 0.2 nM to 8.9 ± 0.8 nM. Again, this increase occurred largely in the nL fraction: 24% to 43% for 7-HDHA, 21% to 45% for 14-HDHA, 28% to 50% for 17-HDHA, and 27% to 48% for 13-HDHA. Overall, supplementation of n3-PUFA resulted in elevated concentrations of EPA and DHA-derived hydroxy-PUFAs in nLs, PC and PE classes, with the strongest increase in nLs (∼600%) (Fig 5 Fig., supplemental Fig. S17). Interestingly, the supplementation led also to a slight increase of ARA-derived hydroxy-PUFA in nLs (Supplemental Fig. S17, Supplemental Table S10, Supplemental Table S11). This lipid class-specific analysis of oxylipins in plasma shows that hydroxy-PUFAs are precursor-dependent esterified to distinct lipid classes and that their lipid class distribution can be shifted with the diet.

**Fig. 5.**
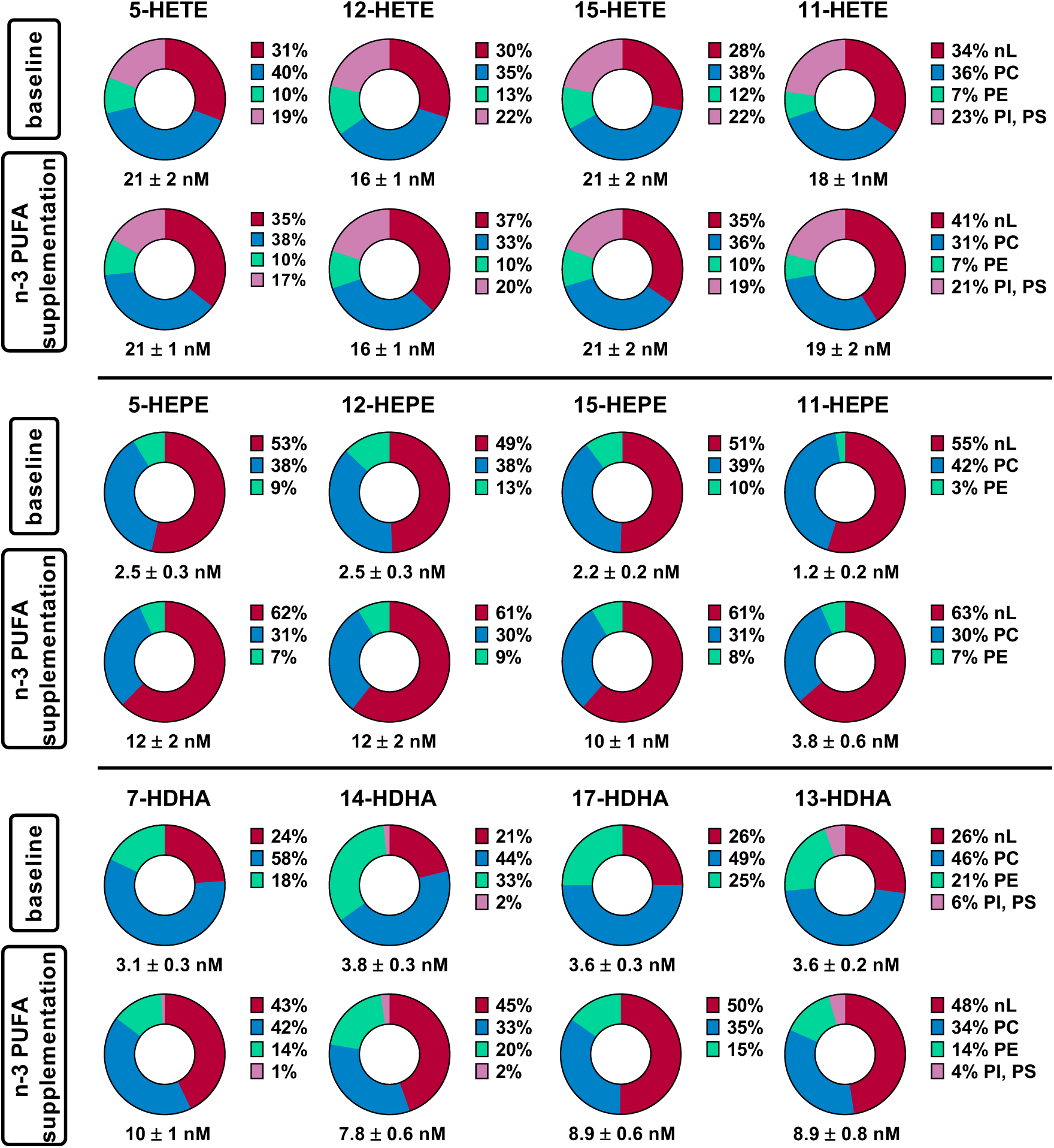
Change in the concentration and distribution pattern of esterified hydroxy-PUFAs across lipid classes in human plasma following 12 months of n3-PUFA supplementation. Total oxylipins were quantified in the lipid class fractions in plasma of human subjects at baseline and after 12 months of n3-PUFA supplementation (1.5 g EPA and 1.8 g DHA per portion; 4 portions per week). Shown is the concentration of total oxylipins and their relative distribution in the lipid classes of a representative set of hydroxy-PUFAs. The concentration of oxylipins in each lipid fraction can be found in Table S10. Analysis was carried out using targeted RP-LC-ESI(-)-MS/MS following alkaline hydrolysis. Shown are the mean values ± SE (n = 9).

## Discussion

A novel lipid class SPE-fractionation method was developed using silica-bonded aminopropyl material as the solid phase. This material has been widely applied in previous lipid fractionation approaches (25, 26, 46). In contrast to commonly used normal-phase LC (NP-LC), our method employed a HILIC-based approach, which has not yet been reported in this context. This approach reduces the use of environmentally hazardous chemicals such as halogenated solvents typically employed in NP-LC. Only a single column is needed to separate the lipids into the classes nLs, PC, PE, PG, and PI/PG with high efficiency. This was shown for the recovery of the spiked labeled lipids in human plasma as well as HEK293 cell extracts. Endogenously present lipids in the samples were also efficiently separated as demonstrated based on lipid class-specific XICs created using MS-DIAL. The chromatographic peak patterns and signal intensities of isolated lipid classes with and without fractionation were similar. Of note, the isolation of the lipid classes led to a strong reduction of ion suppression, improving the LC-MS detection, which could be helpful particularly for the analysis of low-abundant PL classes by untargeted lipidomics.

Compared to SPE-based lipid fractionation protocols previously described, the performance of the separation reported here is better than those using a single column, as they only separated nLs and PLs (47, 48) or did not separate PL classes as effectively, e.g., only isolating PC (49, 50). Methods that have a comparable separation efficiency or which are able to separate five or more lipid fractions employ two or more different SPE cartridges (25–27). Three cartridges were used to separate nLs into MGs, DGs, TGs, and CEs (26, 27) or to separate PLs into PG; PA, PI, PS; PE; as well as LPC, PC and SM (25). Here, we achieved comparable separation efficiencies of the PL classes using only one column, which is ideal for our aim to pinpoint the binding form of esterified oxylipins in major lipid classes. The time needed for sample preparation is thereby short, reducing the risk of artificial formation and degradation, which is a challenge in oxylipin analysis (51).

Quantification of esterified oxylipins in the different lipid fractions after hydrolysis offers better sensitivity compared to direct analysis of intact lipids. Lipid fractionation enabled the detection of hydroxy-PUFAs in PE, PG, and PI/PS in human plasma, which was not reported using a non-targeted (semi-quantitative) LC-HRMS/(MS) analysis (16). Also with a more sensitive targeted LC-MS/MS method on a triple quadrupole instrument, the detection of hydroxy-PUFAs in PLs was not possible in HEK293 cells at baseline (19). This may be explained by the distribution of the oxylipins across a large number of lipid classes and species (Fig. 1). For example, the detection of six hydroxy-PUFAs in the targeted LC-MS/MS oxPL method was based on 63 oxPLs with 190 transitions (three for each oxPL species). The HILIC-based fractionation approach is less specific than direct analyses but provides better sensitivity and covers more oxylipins. Only one chromatographic peak per oxylipin in each lipid fraction results, allowing a sensitive detection. Moreover, the use of authentic oxylipin standards together with a validated LC-MS/MS method (23, 31–34) enables reliable quantification.

The developed fractionation approach was applied to analyze two independent human plasma pools, including NIST SRM 1950 (Supplemental Table S9). The esterified oxylipins present in the samples were quantitatively detected following fractionation because the sum of the concentrations in the lipid fractions matched the concentration of esterified oxylipins determined without fractionation (>80%). Owing to that, this approach allows to quantitatively determine the amount of oxylipins bound in nLs and different PL classes in human plasma. Earlier SPE-based studies only enabled the differentiation of esterified oxylipins between nLs and PLs (29, 52). Watanabe *et al.* investigated the distribution of endogenous epoxy-PUFAs in rat plasma and found that 37% of 11(12)-EpETrE was bound in nLs and 67% in PLs, which is comparable to the distribution of HETEs in human plasma (Fig. 4) (29).

Regarding the binding form of hydroxy-PUFAs in plasma, we found striking precursor-dependent differences (Fig. 4). Oxylipins derived from LA and ALA were preferably bound in nLs. This is consistent with studies investigating oxidized lipids directly. Here, the majority of oxidized LA and ALA was found in nLs (16).

In the Western Diet, which includes high amounts of LA-rich vegetable oils, the consumption of LA (up to 29 g/day) (53) and ALA (up to 2.3 g/day) (54) is much higher compared to the intake of ARA (up to 0.35 g/day) (55) and DHA (up to 0.50 g/day) (56). With that, also considerable amounts of oxylipins are ingested (57–59), which are bound TGs. In plasma, the concentrations of LA-derived oxylipins were many-fold higher compared to oxylipins derived from other PUFAs, consistent with the high consumption of LA and ALA. Because we also found LA and ALA-derived oxylipins in nLs, we assume that these plasma oxylipins may be derived from direct uptake from the diet or from (aut)oxidation of the ingested TGs.

Oxylipins derived from long-chain PUFAs such as ARA and DHA were predominantly esterified to PLs. ARA and DHA are the main PUFAs occurring in cell membranes and are critical for membrane fluidity as well as cell function (60). Moreover, both serve as precursors of bioactive lipid mediators (61, 62). That they, as well as EPA, were found esterified to PLs in plasma suggests that they were released from cells e.g. as vesicles or in lipoproteins.

All analyzed oxylipins showed a distinct distribution among the PL classes. PC is the major PL class in human plasma and contains a large proportion of plasma total FAs (∼ 40%) (43, 63). Thus, it is not surprising that PC is the dominating PL class in which oxylipins are bound in plasma and that all the investigated hydroxy-PUFAs occur at least in part in PC (Fig. 4). Nonetheless, precursor-dependent differences were found: ARA-derived oxylipins and 15-HEPE were present at relevant concentrations in the PI/PS fraction. In cells, ARA is not randomly distributed among PL classes but is preferentially incorporated into PC and PI (64, 65). Here, ARA-PI are one of the precursors for inositol phosphate signaling pathways (66). The occurrence of ARA, its oxylipins and 15-HEPE in PI in plasma may reflect the export of PI from cells, e.g. in the form of vesicles, membrane fragments, or during lipoprotein formation in the liver. While Maskrey *et al.* reported preferential incorporation of 15-HETE into PE in activated monocytes (20) suggesting a direct oxidation of PE by 15-LOX-1 we recently showed that in HEK293 cells, 15-HETE and also 15-HEPE were almost exclusively incorporated into PI via the Lands’ cycle (13, 19). This is consistent with earlier studies using isotopically labeled oxylipins (67, 68). Unlike cell membranes, plasma lipids reflect in addition systemic processes such as lipoprotein metabolism, vesicle transport, or liver-associated export of lipids (69). This may explain why also other ARA-derived oxylipins are found in PI, such as 5-HETE and 12-HETE, which were esterified to PC (and not PI) in HEK293 cells (19).

In plasma DHA and its derived oxylipins showed the highest relative esterification in PE compared to all other lipids. DHA is known to be preferentially enriched in PE in the inner leaflet of membranes, e.g. as a regulator of membrane fluidity in human cells (70). The release of oxPE would thus explain the high concentration of DHA and its derived oxylipins in the PE fraction. Of note, consistent with our studies in HEK293 cells (13, 19), DHA-derived hydroxy-PUFAs do not occur in PI in human plasma.

We then asked the question whether n3-PUFA supplementation changes the oxylipin binding forms in human plasma. It is well described that the composition of non-esterified (71–75) and esterified oxylipins (14, 76–78) in plasma is dramatically changed by n3-PUFA supplementation. At baseline, the participants showed the same distribution of HETEs, HEPEs, and HDHAs over the lipid classes as the two other analyzed plasma pools (Supplemental Fig. S13 – S15). Consistent with earlier studies (74, 75), n3-PUFA supplementation hardly changed the concentration of HETEs while oxylipins derived from HEPEs and HDHAs increased dramatically (Fig. 4). EPA showed a greater relative increase due to its lower baseline level (76, 78) (Supplementation Fig. S16). This increase in plasma concentrations of n3-PUFA-derived oxylipins by supplementation can be explained in different ways. First, the oxylipins are present in the (fish oil) n3-PUFA supplements due to autoxidation (79), which are then absorbed together with n3-PUFAs and released in the bloodstream. Ostermann *et al.* reported a perfect dose-dependent increase of n3-PUFA-derived non-esterified oxylipins following n3-PUFA supplementation in plasma (76), which could be well explained by the co-ingestion and absorption of HEPEs and HDHAs with their precursor PUFAs, EPA and DHA, respectively. The second explanation is that the n3-PUFA are first absorbed, integrated into lipids, and the oxylipins are then formed by autoxidation or enzymatically, e.g., by LOX (6). Oxidation of PUFAs can occur with or without release by lipase and re-esterification by Lands’ cycle enzymes (80). Because LOX products such as 15-HEPE or 17-HDHA are similarly elevated as autoxidatively formed 11-HEPE and 13-HDHA, enzymatic formation may not be the dominating factor.

Although supplementation increased the concentration of HEPEs and HDHAs in all lipid classes, the highest elevation was surprisingly found in nLs. This distinct increase supports a direct absorption of oxylipins alongside the n3-PUFAs, similarly as discussed for LA and ALA. The biological roles of esterified oxylipins in plasma remain poorly understood. Shearer and Newman suggest that esterification serves as a mechanism to regulate the biological activity of oxylipins (81). Based on that, the elevated intake/formation of PUFAs and oxylipins following supplementation could thus result in an increased esterification into nLs to control the bioactivity of the oxylipins functioning as a reservoir. The esterified oxylipins can be released from lipoproteins such as VLDL by lipoprotein lipases and then may act on endothelial cells (71, 81, 82). Our data shows that even after 12 months supplementation, the DHA-derived hydroxy-PUFAs occur in plasma more in nLs compared to baseline, suggesting a direct effect of the diet on the plasma oxylipins. Many studies aim to correlate plasma oxylipin levels with health status, ideally serving as biomarkers for diseases (83–85). The high variability in plasma oxylipins observed in these studies may be explained by direct effects from the diet, which warrant further investigation.

## Conclusion

This study provides novel insights into the lipid class-specific occurrence of oxylipins in human plasma. A new HILIC-based fractionation method enabled the efficient separation of lipids into nLs and four PL fractions using a single SPE cartridge. This approach allows the quantification of a large number of esterified oxylipins in lipid fractions and can readily be integrated with existing oxylipin analysis workflows. The data demonstrate in two independent pools of human plasma that esterification patterns of oxylipins in plasma are largely determined by their precursor PUFAs. Oxylipins derived from LA and ALA are largely bound to nLs, presumably reflecting the circulation of oxylipins which have been ingested with the diet alongside their precursor PUFAs. In contrast, oxylipins derived from the long-chain PUFAs ARA, EPA and DHA circulate in plasma primarily esterified in PLs, presumably reflecting their release from cellular membranes, e.g. as lipoproteins or vesicles. Moreover, the preferred esterification of ARA, its oxylipins and 15-HEPE to PI and of DHA and its oxylipins to PE in human cells is reflected in plasma lipids. Notably, n3-PUFA supplementation resulted in an overall increase of HEPEs and HDHAs combined with a significant redistribution of HDHAs toward binding in nLs. This raises new questions regarding the direct effect of diet on the occurrence of esterified oxylipins in human plasma.

## Abbreviations

ACN: acetonitrile
AGC: active gain control
ALA: α-linolenic acid (18:3(9Z,12Z,15Z))
ARA: arachidonic acid (20:4(5Z,8Z,11Z,14Z))
BHT: butylhydroxytoluene
Cer: ceramide
Chol Ester: cholesterol ester
COX: cyclooxygenase
CYP: cytochrome P450 monooxygenase
dd: data-dependent
DG: diacylglycerol
HDHA: hydroxydocosahexaenoic acid
HEPE: hydroxyeicosapentaenoic acid
HILIC: hydr^1^ophilic interaction liquid chromatography
HRMS: high resolution mass spectrometry
IPA: isopropanol
IS: internal standard
LA: linoleic acid (18:2(9Z,12Z))
LC: liquid chromatography
LLOQ: lower limit of quantification
LOX: lipoxygenase
Lyso-PC: lyso-phosphatidylcholine
Lyso-PE: lyso-phosphatidylethanolamine
MeOH: methanol
MG: monoacylglycerol
NIST: US National Institute of Standards and Technology
nL: neutral lipid
NP: normal-phase
PC: phosphatidylcholine
PE: phosphatidylethanolamine
PE-O: ether phosphatidylethanolamine
PG: phosphatidylglycerol
PI: phosphatidylinositol
PL: phospholipid
PS: phosphatidylserine
RP: reversed-phase
SPE: solid-phase extraction
SPLASH: Single-vial Prepared Lipidomic Analytical Standard for Human plasma lipids
t-AUCB: trans-4-[4-(3-adamantan-1-yl-ureido)-cyclohexyloxy]-benzoic acid
XIC: extracted ion chromatogram

## Declarations

Blood for the n3-PUFA supplementation study was collected from healthy human subjects in accordance with the guidelines of the Declaration of Helsinki and approved by the ethics committee of the University of Wuppertal. The n3-PUFA supplementation study was approved by the Suffolk Local Research Ethics Committee.

### Data availability

All data are contained within the article or supplementary material.

## Supplemental Data

This article contains Supplemental data (13, 30, 32, 33, 36, 47, 86–88)

## Supplemental data

**Fig S1:**
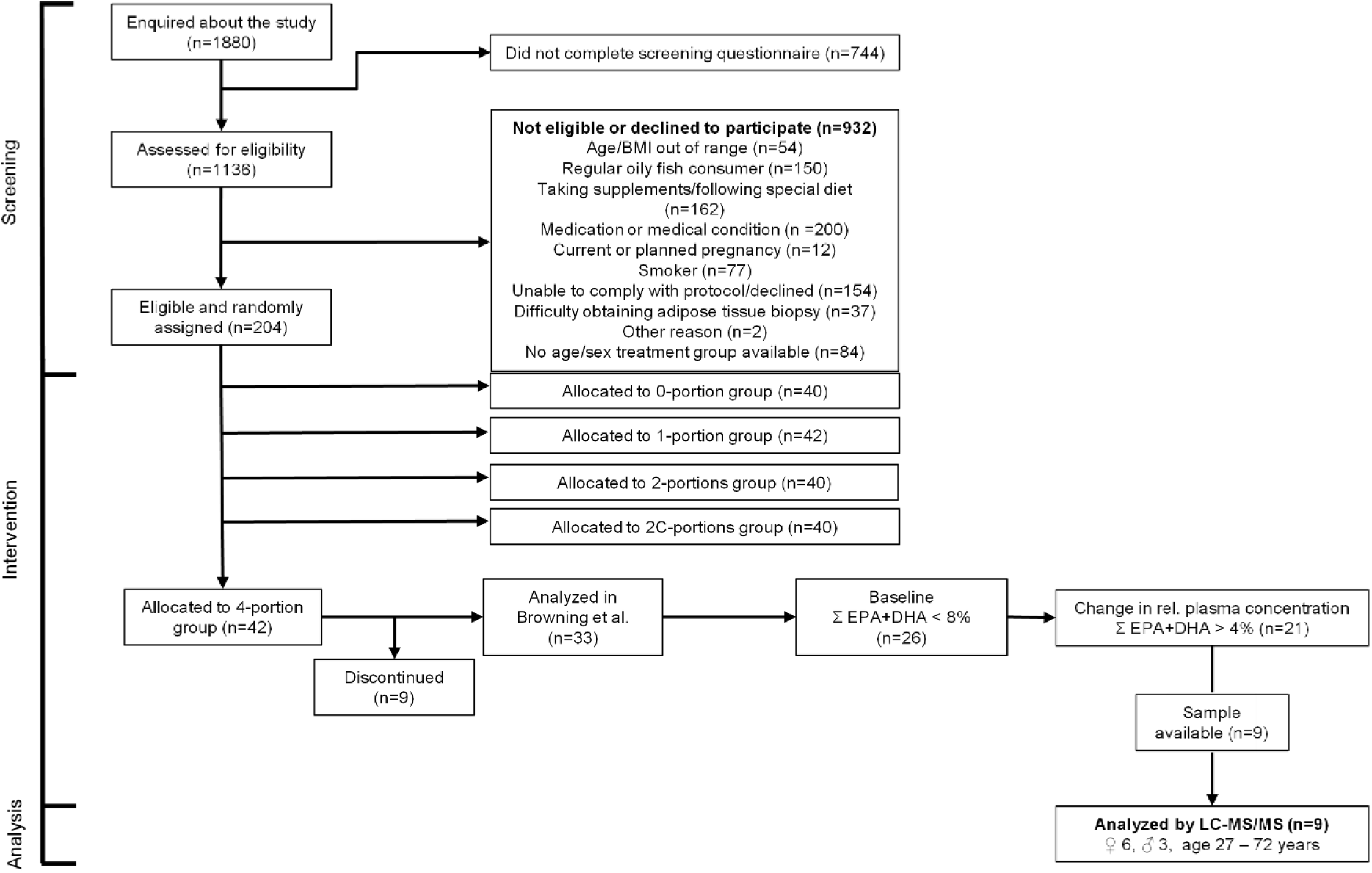
Design of the n3-PUFA supplementation study and sample selection. A subset of human plasma samples from a clinical study (1, 2) was used to investigate the pattern of esterified oxylipins in different lipid classes following n3-PUFA supplementation. 9 participants (3 males, 6 females, age 23 – 72 years) out of 42 subjects were selected fulfilling the following criteria: On 4 days per week subjects received n3-PUFA capsules containing 1.5 g EPA and 1.8 g DHA per portion, corresponding to 4 portions of fatty fish a week (4-portion group). Plasma samples had a relative level of EPA + DHA <8% of total FA at baseline and a change in the relative level of EPA + DHA >4% after 12 months of n3-PUFA supplementation (1). Available aliquots of the samples at baseline and after 12 months were properly stored at -80°C. Plasma samples were fractionated and esterified FA and oxylipins were analyzed after alkaline hydrolysis by LC-ESI(-)-MS/MS (3–5).

**Fig. S2:**
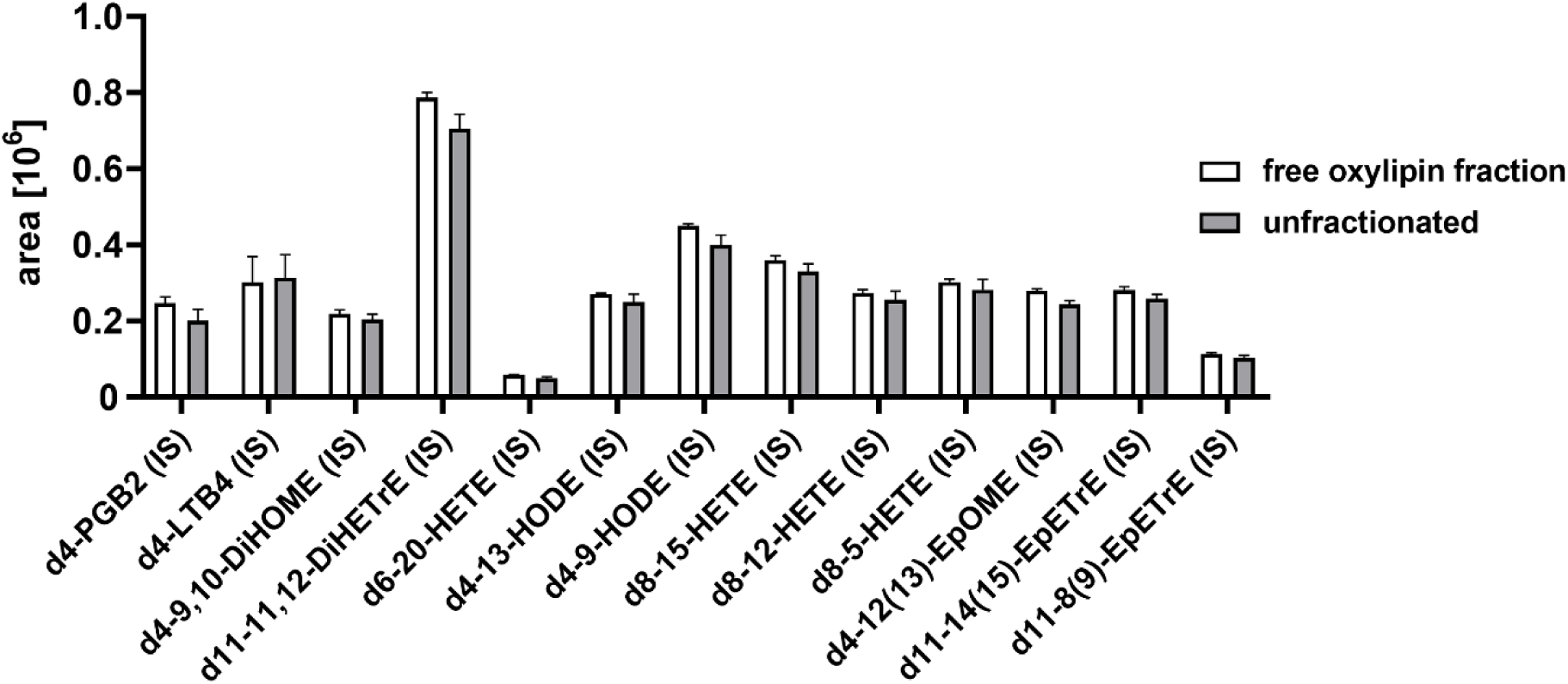
Recovery of free oxylipins following SPE fractionation of lipid classes. Human plasma (100 µL) was spiked with a mixture of deuterium-labeled oxylipins followed by fractionation of lipid classes using SPE. The fraction containing free oxylipins was collected and free oxylipins were analyzed. The peak areas of the standards in the fraction were compared to those of the standard mixture directly analyzed without fractionation. The comparison of the peak areas shows a high recovery of free oxylipins after SPE indicating that free oxylipins are completely eluted by 18 mL MeOH with 0.1% acetic acid (Fig. 2) Analysis of oxylipins was carried out using targeted LC-ESI(-)MS/MS (3–5). Results are shown as mean ± SD (n = 3).

**Fig. S3:**
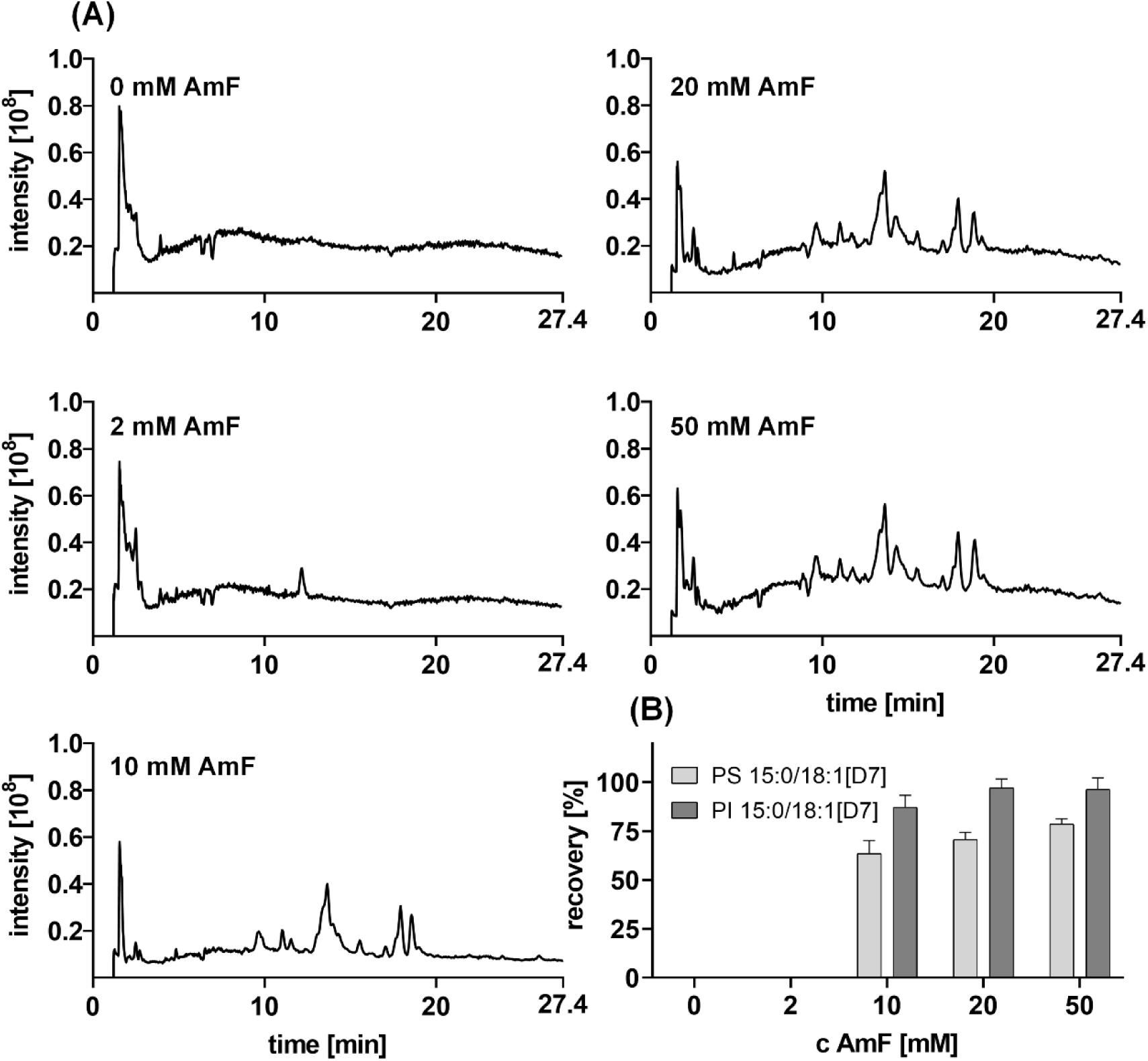
Ammonium formate-dependent elution of PI and PS species in lipid fraction 5. **(A)** Shown are total ion chromatograms (*m/z* 200 - 1200) of SPE lipid fraction 5 of a HEK293 cell extract eluted separately with increasing concentrations of ammonium formate (AmF). The eluent was composed of H_2_O/ACN/IPA (20:35:45, v/v/v) containing 0.1% formic acid and ammonium formate (0 – 50 mM). **(B)** Recovery of PI 15:0/18:1[D7] and PS 15:0/18:1[D7] with increasing ammonium formate concentration used for elution (n = 3). The results indicate that a minimum ammonium formate concentration of 10 mM in the eluent is required for effective elution of PI and PS species; 20 mM was chosen. Analysis was carried out using untargeted LC-ESI(-)-HRMS (Q Exactive HF) in Full MS/ddMS^2^ TOP N mode (6)

**Fig. S4:**
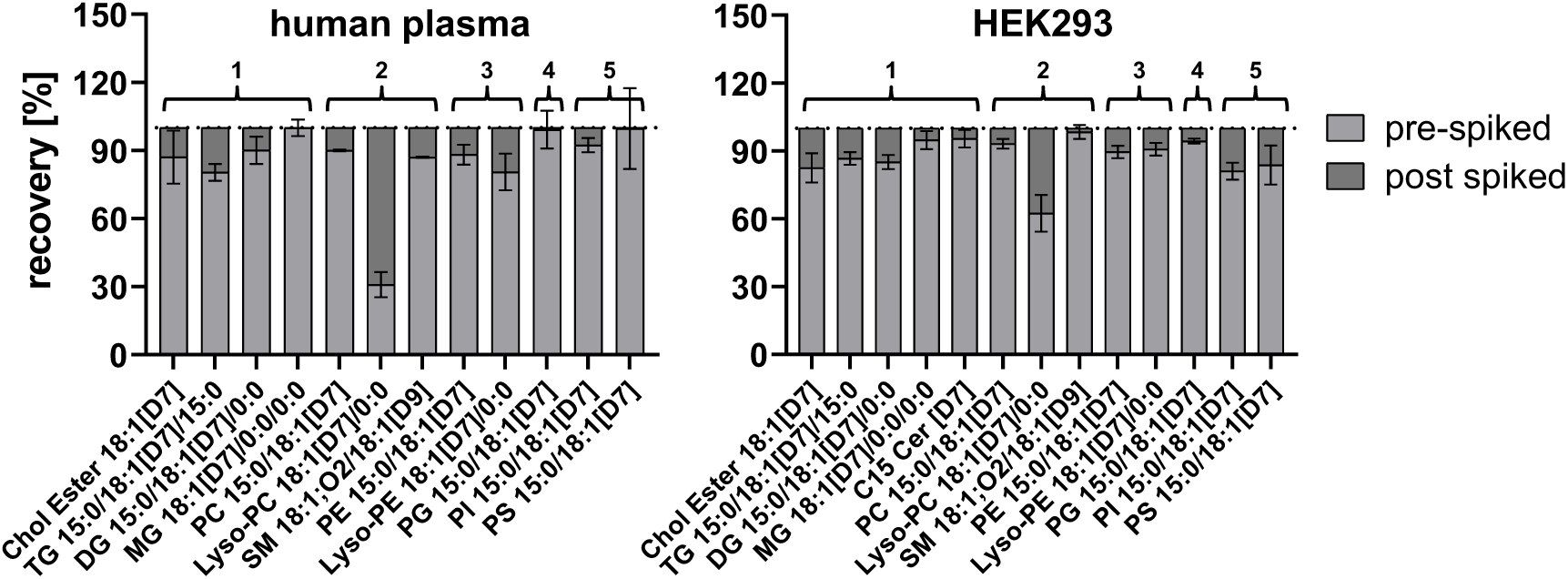
Recovery of deuterium-labeled lipids (SPLASH standards) after lipid class fractionation using SPE. Human plasma (100 µL) or HEK293 cell homogenate (⁓400 µg protein) were extracted and spiked with SPLASH Lipidomix or EQUI SPLASH, and samples were fractionated using SPE. Another set of both samples was spiked with the standards after SPE-fractionation in each individual fraction (post-spiked), which was set to 100% recovery. The results show high extraction recoveries of most standards after fractionation, with more than 80% recovered in each fraction. Only lyso-PC 18:1[D7]/0:0 shows lower recovery in fraction 2 because it partially coelutes in fraction 3. The number of the lipid fractions is shown at the top of the figure. Analysis was carried out using untargeted LC-ESI-HRMS (Q Exactive HF) in Full MS/ddMS^2^ TOP N mode (6). Results are shown as mean ± SD (n = 3).

**Fig. S5:**
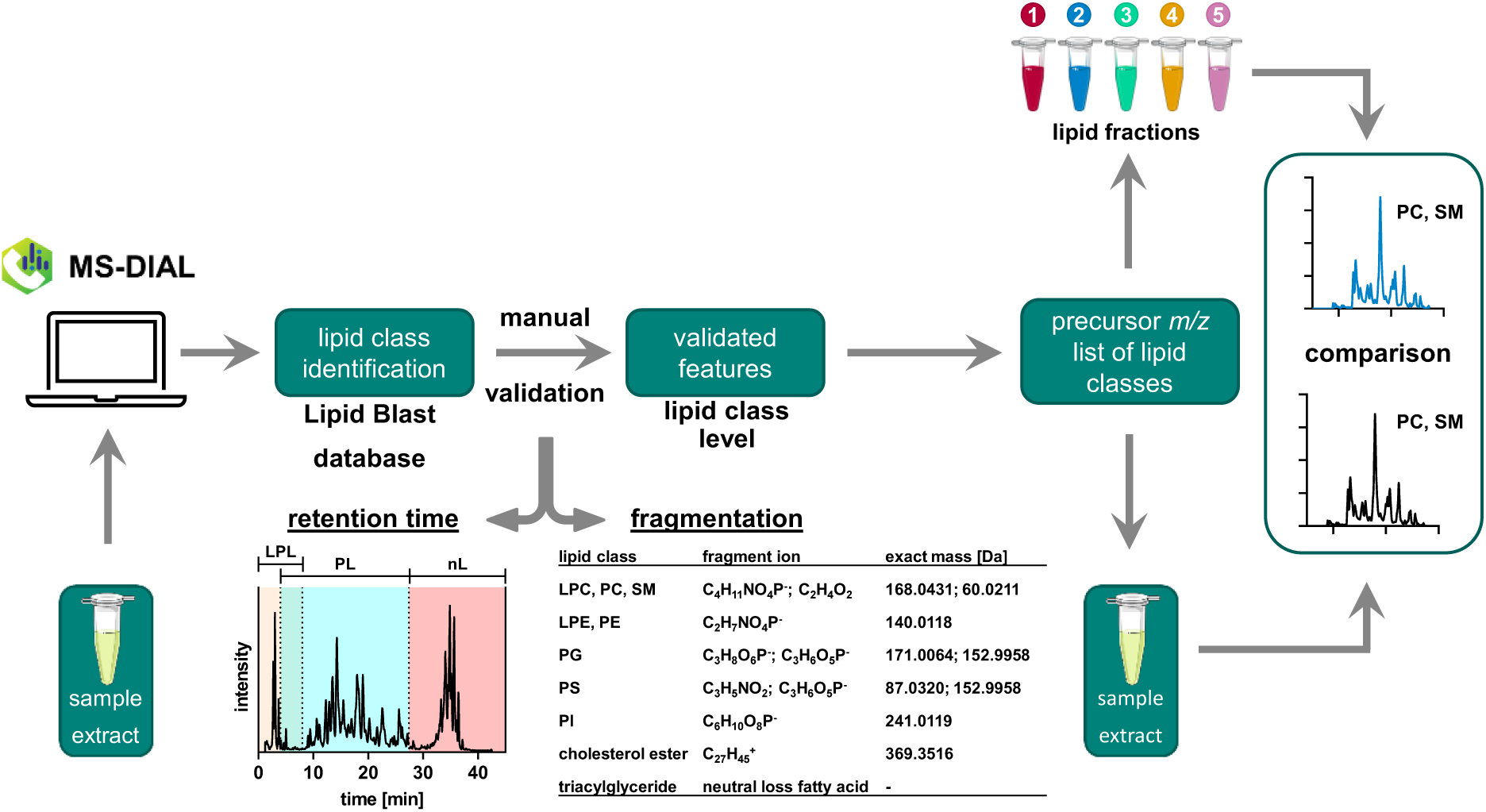
Data processing for chromatographic lipid class pattern comparison. Lipids were extracted and analyzed by untargeted LC-HRMS (Q Exactive HF Orbitrap) in Full MS/ddMS² TOP N mode (Table S7). Data evaluation was carried out in both ionization modes using MS-DIAL software (version 5.3) (7). The processing steps included feature detection, spectra deconvolution, and peak alignment across samples. Detailed data processing and parameter settings for MS-DIAL are listed in Table S6. For evaluation, only features matching the used spectral library were used (reference matched) (Tables S2 and S4). The features were manually validated for lipid class identification according to Liebisch *et al*. (8) based on plausible retention times and fragmentation criteria (Table S3 and Table S5). The retention time criteria for validating the features were as follows: lyso-PL (≤8 min), PL (≥4 min), SM (≥4 min), CE, and TG (≥27.4 min). Features outside these criteria were excluded from further evaluation. As fragmentation criteria, characteristic product ions of the lipid headgroup were used, such as *m/z* 168.0426 for PC and SM, *m/z* 140.0118 for PE, or *m/z* 241.0113 for PI in negative mode according to Pi *et al.* (9) (Fig. S9-S12). Validated features (lipid class level) were listed and sorted according to the SPE-based lipid class fractionation (Tables S2, S3). Theoretical precursor *m/z* values assigned to lipids eluting in the corresponding fraction were used to generate summed XICs for all lipids of the eluting lipid classes. XICs were generated for both non-fractionated samples and fractionated samples to assess lipid recovery within specific lipid classes after fractionation. Analysis was carried out using untargeted LC-ESI-HRMS (Q Exactive HF) in Full MS/ddMS^2^ TOP N mode (6) with slight modifications. The following gradient was used: 0–0.7 min 30% B; 0.7–0.8 min 30–52.5% B; 1.5–11 min 52.5% B; 11–20 min 52.5–60% B; 20–40 min 60–99% B; 40–42 min 99% B; 42–45 min 30% B with a flow rate of 260 µL/min. The method included a polarity switch to detect different lipids in both negative and positive ionization modes within a single run. The analysis started in ESI(-) mode to detect and characterize polar lipids, such as PLs, and was switched to ESI(+) mode at 27.4 min to detect nL, such as TGs and CEs. Detailed instrument parameters are given in Table S7.

**Fig. S6:**
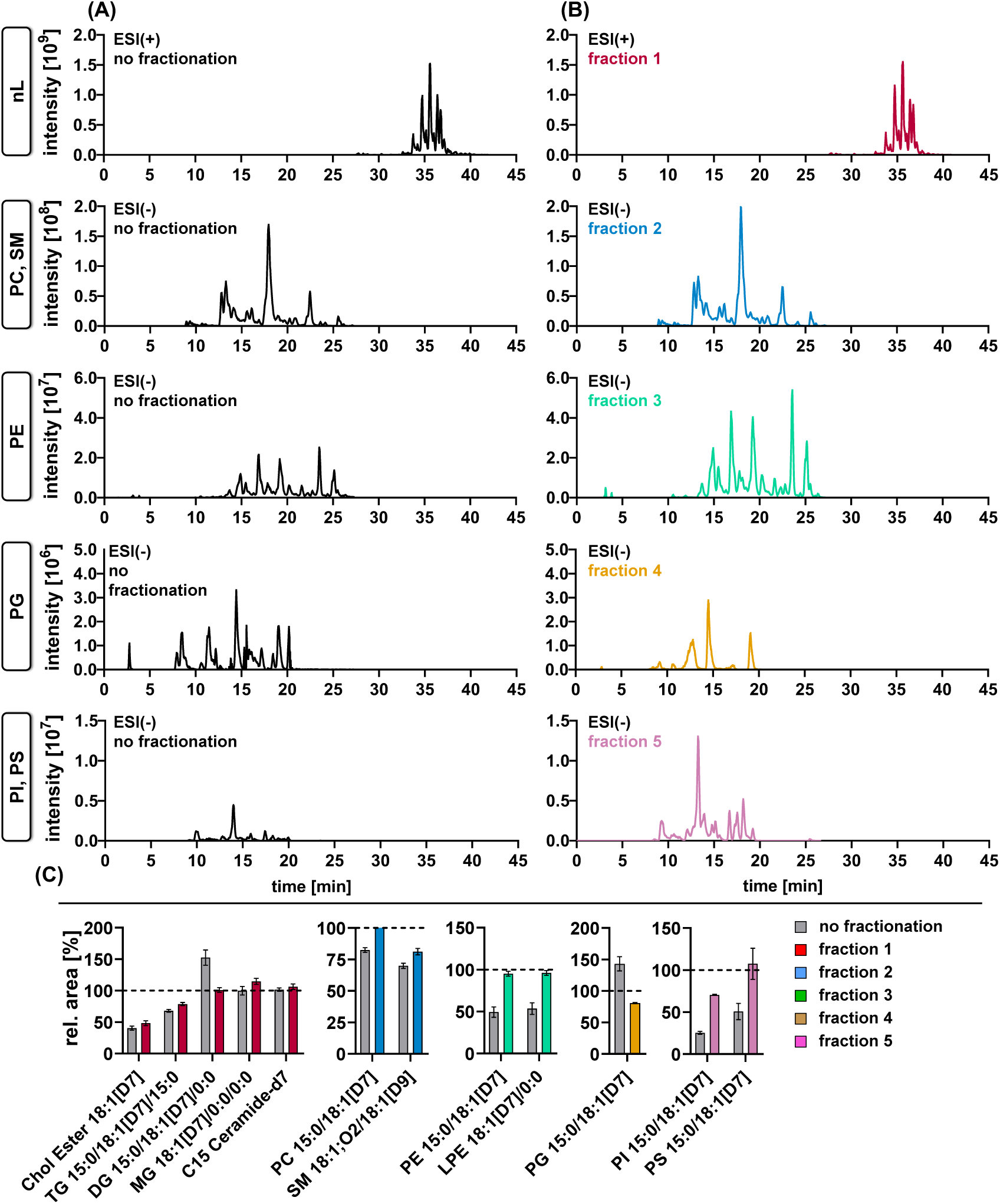
SPE fractionation of lipid classes in HEK293 cells. Shown are cumulated XICs (sum of all detected species, Fig. S5) of nLs and different PL classes in HEK293 cells **(A)** without and **(B)** with fractionation. XICs were defined based on validated lipid features using MS-DIAL (see Table S5). Theoretical precursor m/z of identified and validated features on lipid class level were used to build XICs in the non-fractionated and the fractionated samples according to the separated lipid classes in each fraction (see Fig S4 for details). **(C)** Recovery of IS in the fractions compared to the non-fractionated sample shows a reduction of ion suppression effects for PE, PG, and PI/PS.

**Fig. S7:**
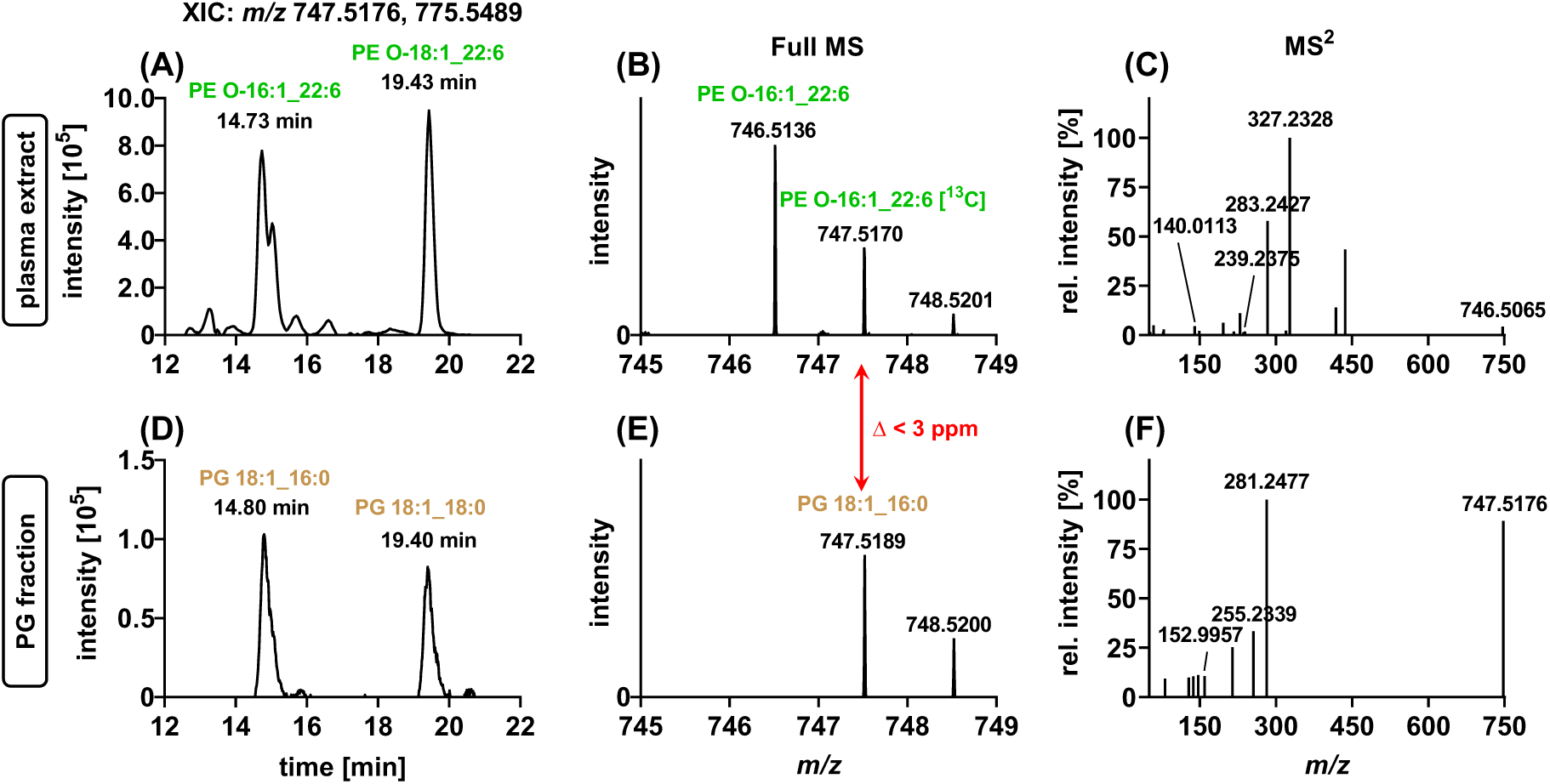
Co-elution of PG and PE-O makes the detection of PG in biological samples by RP-LC-HRMS impossible, which can be solved by lipid class fractionation. **(A)** XIC of *m/z* 747.5182 and *m/z* 775.5443 (≤5 ppm) corresponding to the monoisotopic *m/z* of characterized PE O-16:1_22:6 and PE O-18:1_22:6 in a non-fractionated human plasma extract. **(B)** Zoomed Full MS spectrum at 14.73 min and **(C)** ddMS^2^ scan of the precursor *m/z* 746.5136 (isolation window *m/z* 1.5). **(D)** XIC of *m/z* 747.5176 and *m/z* 775.5489 (≤5 ppm) corresponding to characterized PG species PG 18:1_16:0 and PG 18:1_18:0 in SPE fraction 4. **(E)** Zoomed Full MS and **(F)** ddMS^2^ scan of *m/z* 747.5189 (isolation window ± *m/z* 1.5). **(A)** shows that co-eluting and higher abundant PE O-species overlay PG species in human plasma. The ions of the M+1 isotope of PE O-16:1_22:6 and PE O-18:1_18:0 are nearly isobaric (≤ 5 ppm), respectively, with PG 18:1_16:0 and PG 18:1_18:0 making it impossible (for MS-DIAL) to detect and characterize the PG species in the non-fractionated human plasma extract. The results show that fractionation can provide increased confidence in determining the lipid class of a lipid, as well as facilitating the detection and characterization of less abundant lipids that coelute with abundant isobaric lipids. Analysis was carried out using untargeted LC-ESI-HRMS (Q Exactive HF) in Full MS/ddMS^2^ TOP N mode (6) with modifications described in Fig. S5.

**Fig S8:**
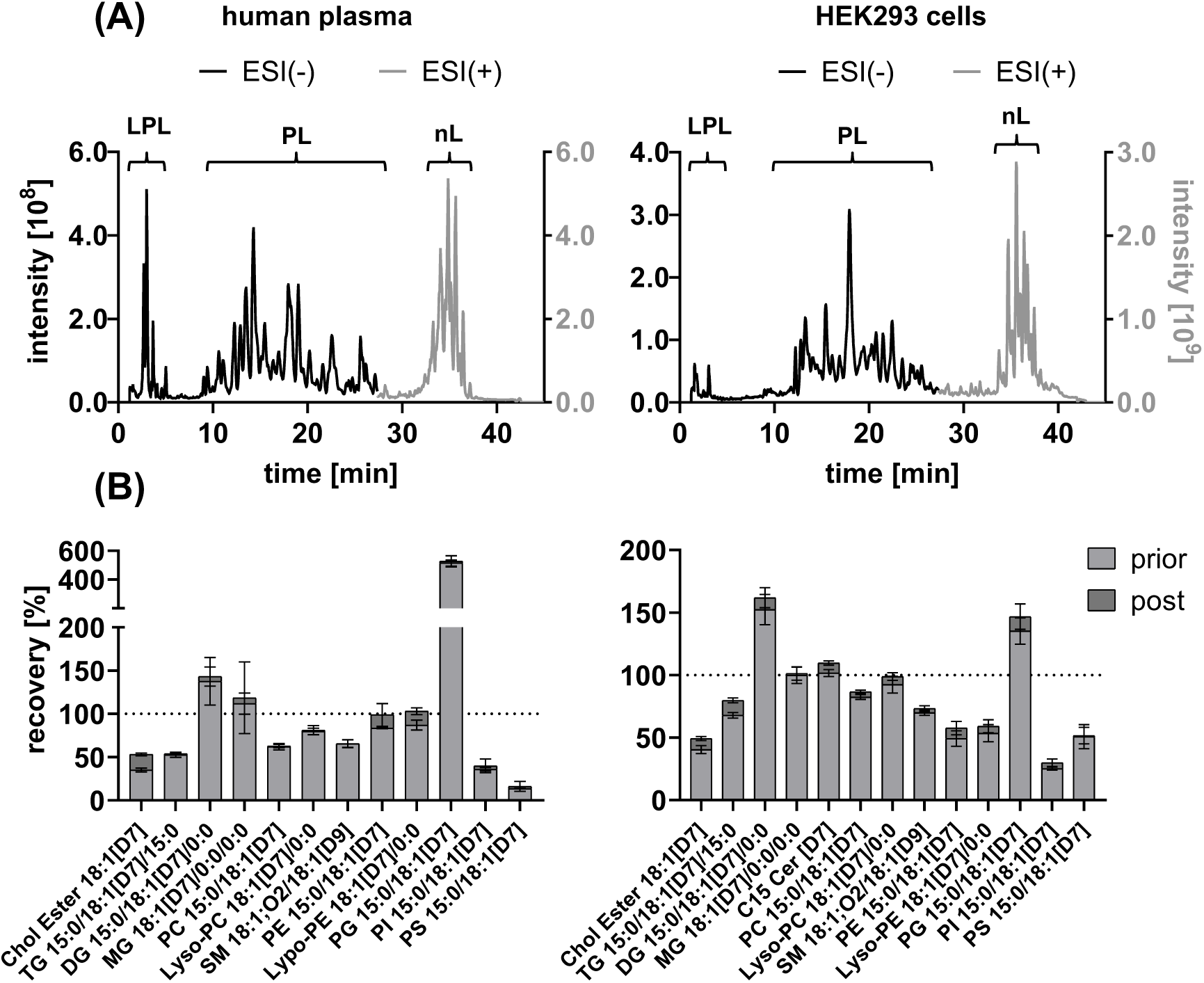
Analysis of human plasma and HEK293 cells using untargeted LC-HRMS. **(A)** Total ion chromatograms of extracts from human plasma and HEK293 cells. (B) Recovery of ISs (SPLASH Lipidomics for human plasma and Equi-SPLASH for HEK293T cells) after extraction. Samples were spiked with IS both either before (prior) or after (post) extraction, and IS peak areas were compared. The results indicate high extraction recovery (>80%) for most lipids. Severe matrix effects were observed, particularly for PI 15:0/18:1[D7] and PS 15:0/18:1[D7] in both, human plasma and HEK293 cells. Figures 2 and S5 demonstrate that these suppression effects can be mitigated by SPE-fractionation of lipids. Analysis was carried out using untargeted LC-ESI-HRMS (Q Exactive HF) in Full MS/ddMS2 TOP N mode (6), with modifications described in Fig. S5. Results are shown as mean ± SD (n = 3).

**Fig. S9:**
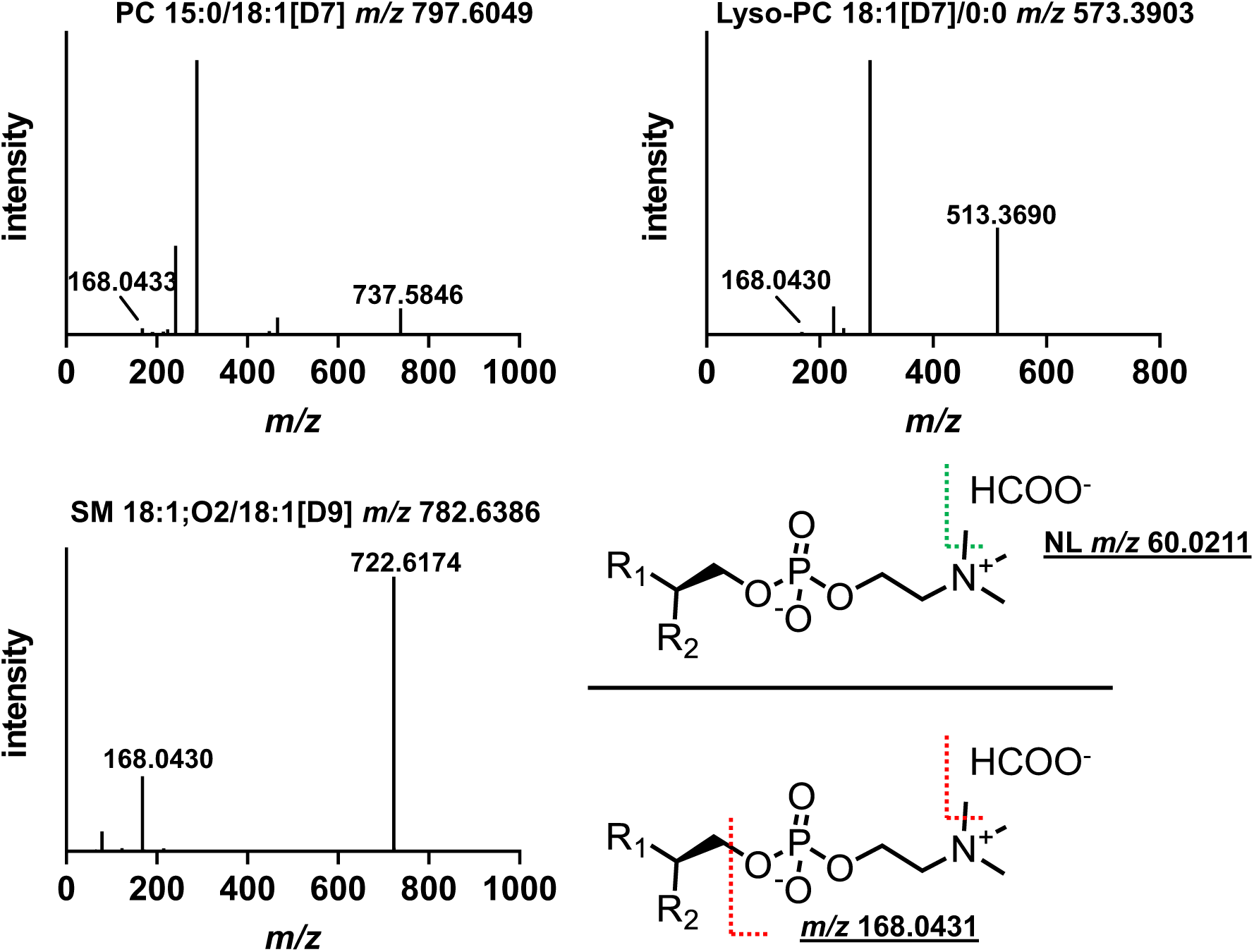
Characteristic fragment ions of choline-bearing phospholipids in LC-ESI(-)MS/MS. Shown are MS^2^ spectra of [M+HCOO^-^]^-^ ions of PC 15:0/18:1[D7], lyso-PC 15:0/18:1[D7] and SM 18:1;O2/18:1[D9] in ESI(-) mode. The spectra show a similar fragmentation behavior of the choline headgroup for all the compounds: 1) neutral loss (NL) of *m/z* 60.0211 by demethylation and loss of the formate adduct and 2) demethylated [M-H]^-^ choline phosphate fragment at *m/z* 168.0431 in line with Pi et al. (9). Analysis was carried out using untargeted LC-ESI(-)HRMS (Q Exactive HF Orbitrap) in Full MS/ddMS² mode (6) with modifications described in Fig. S5.

**Fig. S10:**
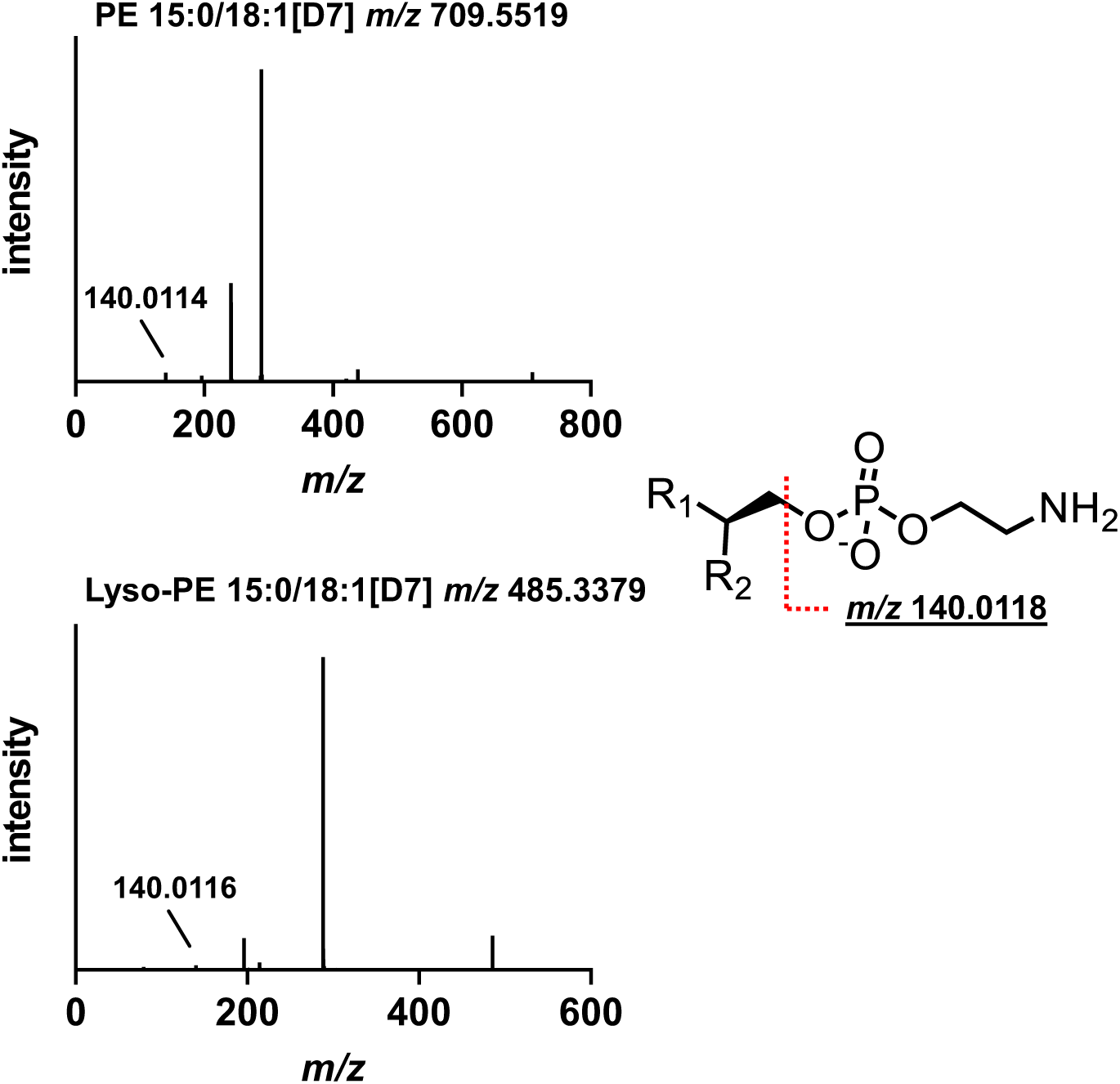
Characteristic fragment ions of ethanolamine-bearing phospholipids in LC-ESI(-)MS/MS. Shown are MS^2^ spectra of [M-H]^-^ ions of PE 15:0/18:1[D7] and lyso-PE 15:0/18:1[D7] in ESI(-) mode. The spectra show similar suggested fragmentation behavior of the ethanolamine headgroup for both compounds: ethanolamine phosphate fragment [M-H]^-^ at *m/z* 140.0118 in line with Pi et al. (9). Analysis was carried out using untargeted LC-HRMS (Q Exactive HF Orbitrap) in Full MS/ddMS² mode (6) with modifications described in Fig. S5.

**Fig. S11:**
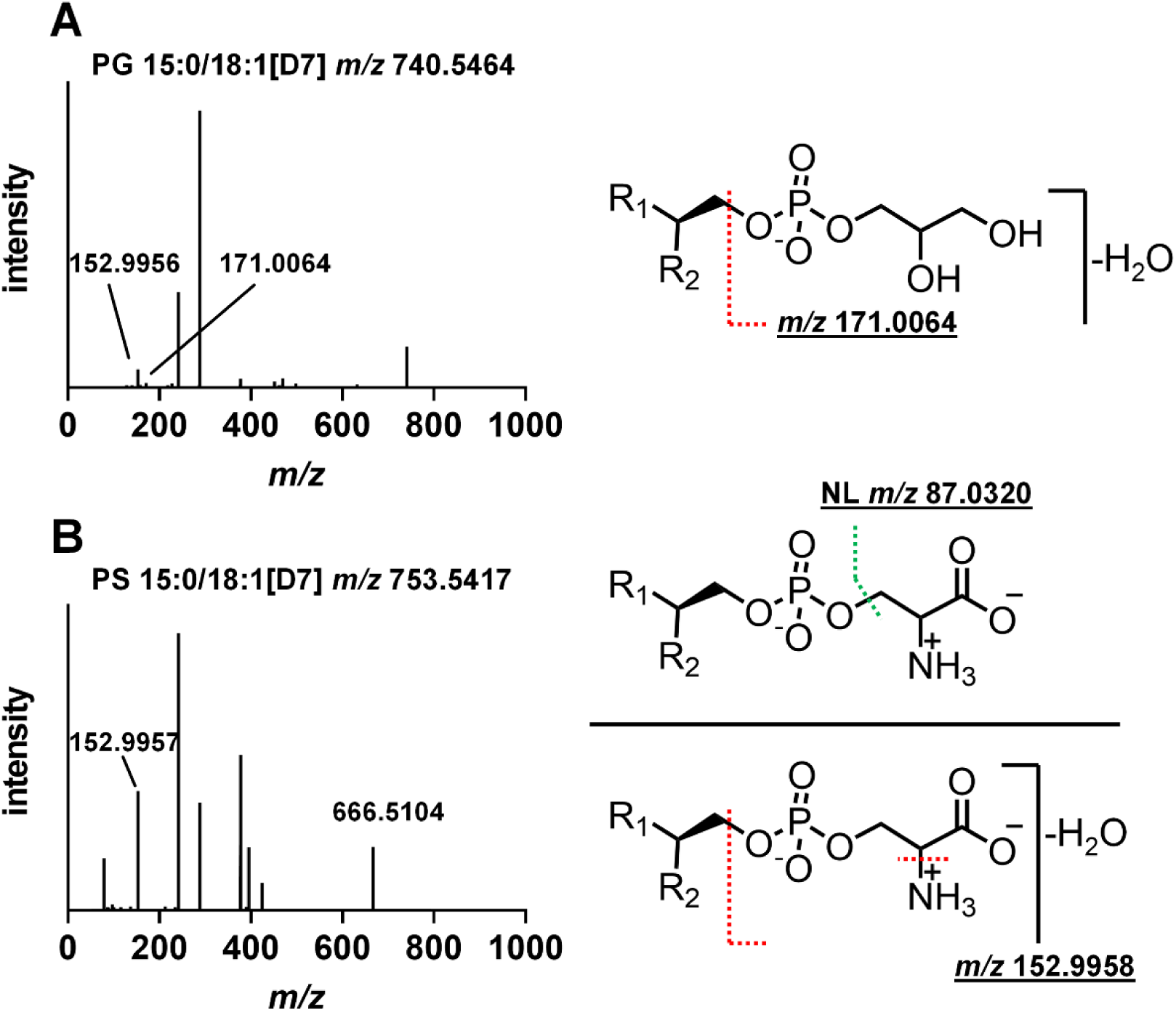
Characteristic fragment ions of glycerol- and serine-bearing phospholipids in LC-ESI(-)MS/MS. **(A)** MS^2^ spectrum of PG 15:0/18:1[D7] [M-H]^-^ in ESI(-) mode. The spectrum shows a characteristic fragment ion of PG: glycerol phosphate fragment [M-H]^-^ with H_2_O loss at *m/z* 171.0064. **(B)** MS^2^ spectrum of PS 15:0/18:1[D7] [M-H]^-^ in ESI(-) mode. The spectrum shows characteristic fragment ions of PS: neutral loss of *m/z* 87.0320 through loss of the serine group in combination with the serine phosphate fragment [M-H]^-^ after deamination and loss of H_2_O at *m/z* 152.9958 in line with Pi et al. (9). Analysis was carried out using untargeted LC-HRMS (Q Exactive HF Orbitrap) in Full MS/ddMS² mode (6) with modifications described in Fig. S5.

**Fig. S12:**
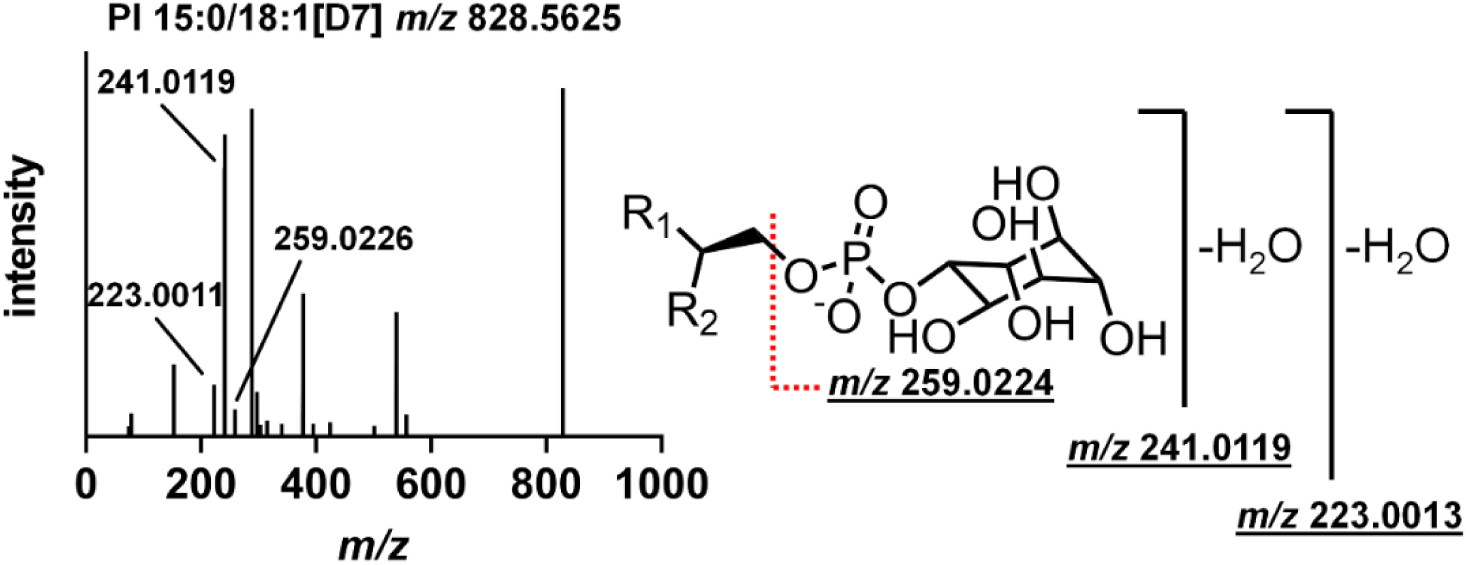
Characteristic fragment ions of inositol-bearing phospholipids in LC-ESI(-)MS/MS. Shown is a MS^2^ spectrum of [M-H]^-^ ion of PI 15:0/18:1[D7] in ESI(-) mode. The spectrum shows characteristic fragment ions of PI: inositol phosphate fragment [M-H]^-^ at *m/z* 259.0224 with loss of one H_2_O at *m/z* 241.0119 and loss of two H_2_O at *m/z* 223.0013 in line with Pi et al. (9). Analysis was carried out using untargeted LC-HRMS (Q Exactive HF Orbitrap) in Full MS/ddMS² TOP N mode (6) with modifications described in Fig. S5.

**Fig. S13.**
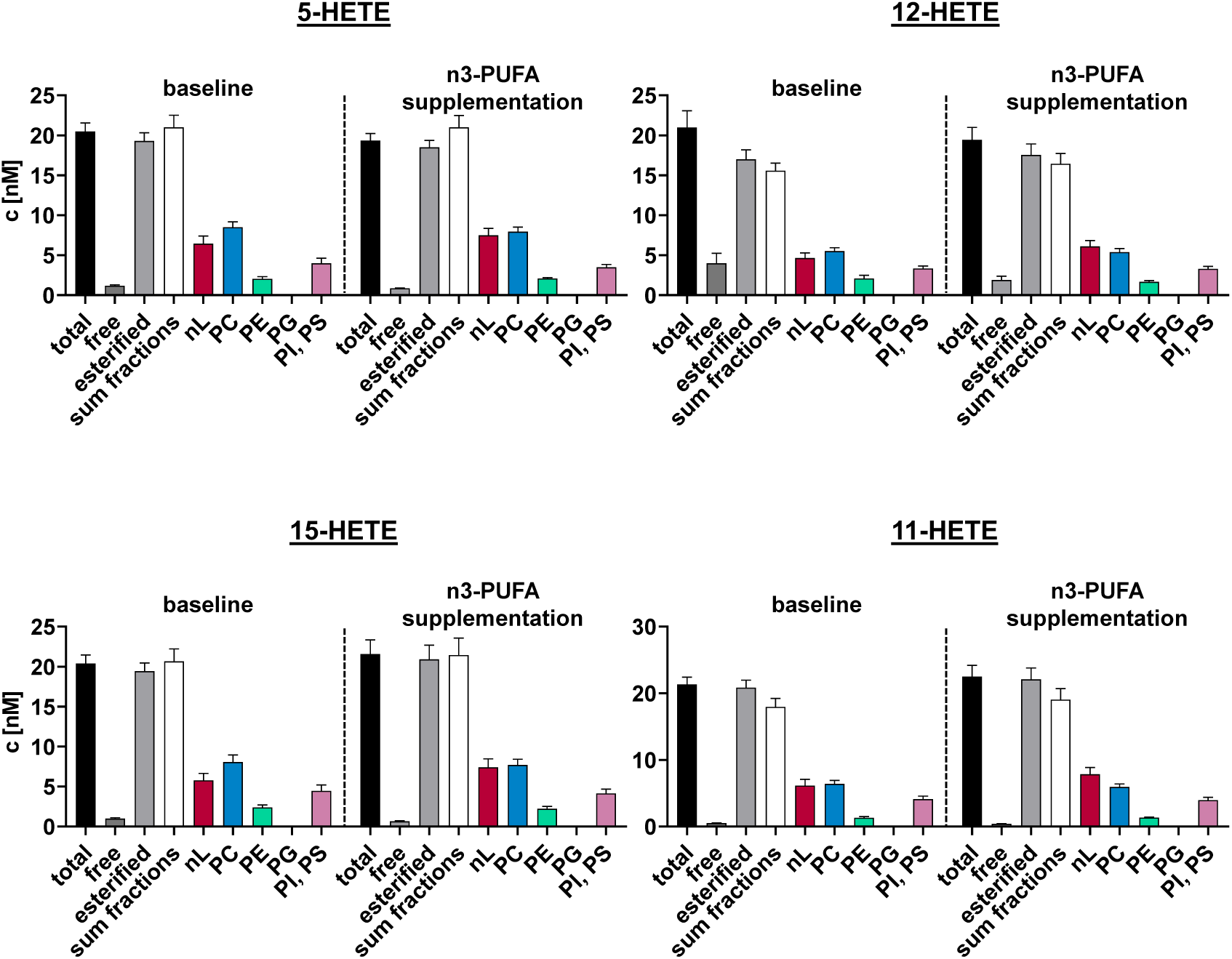
Concentration and distribution of hydroxy-ARA in human plasma esterified in different lipid classes at baseline and after 12 months of n3-PUFA supplementation. Total oxylipins were quantified in the lipid class fractions in plasma of human subjects at baseline and after 12 months of n3-PUFA (1.5 g EPA and 1.8 g DHA/portion; 4 portions per week) supplementation (n = 9). Shown is the concentration ± SE (n = 9) of free and total hydroxy-ARA. All determined concentrations can be found in Table S10. Analysis was carried out using targeted RP-LC-ESI(-)- MS/MS (3–5).

**Fig. S14.**
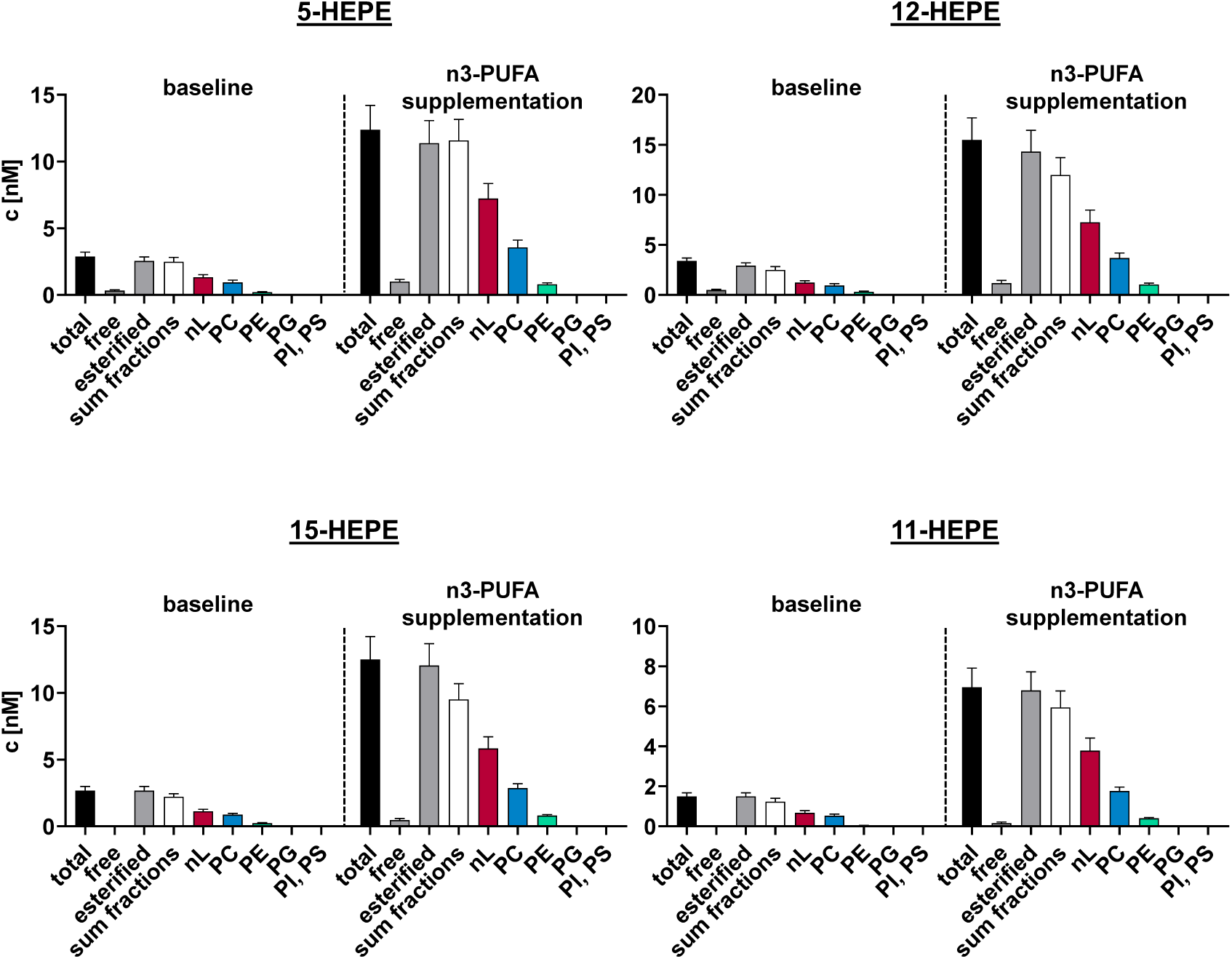
Concentration and distribution of hydroxy-EPA in human plasma esterified in different lipid classes at baseline and after 12 months of n3-PUFA supplementation. Total oxylipins were quantified in the lipid class fractions in plasma of human subjects at baseline and after 12 months of n3-PUFA (1.5 g EPA and 1.8 g DHA/portion; 4 portions per week) supplementation (n = 9). Shown is the concentration ± SE (n = 9) of free and total hydroxy-EPA. All determined concentrations can be found in Table S10. Analysis was carried out using targeted RP-LC-ESI(-)- MS/MS (3–5).

**Fig. S15.**
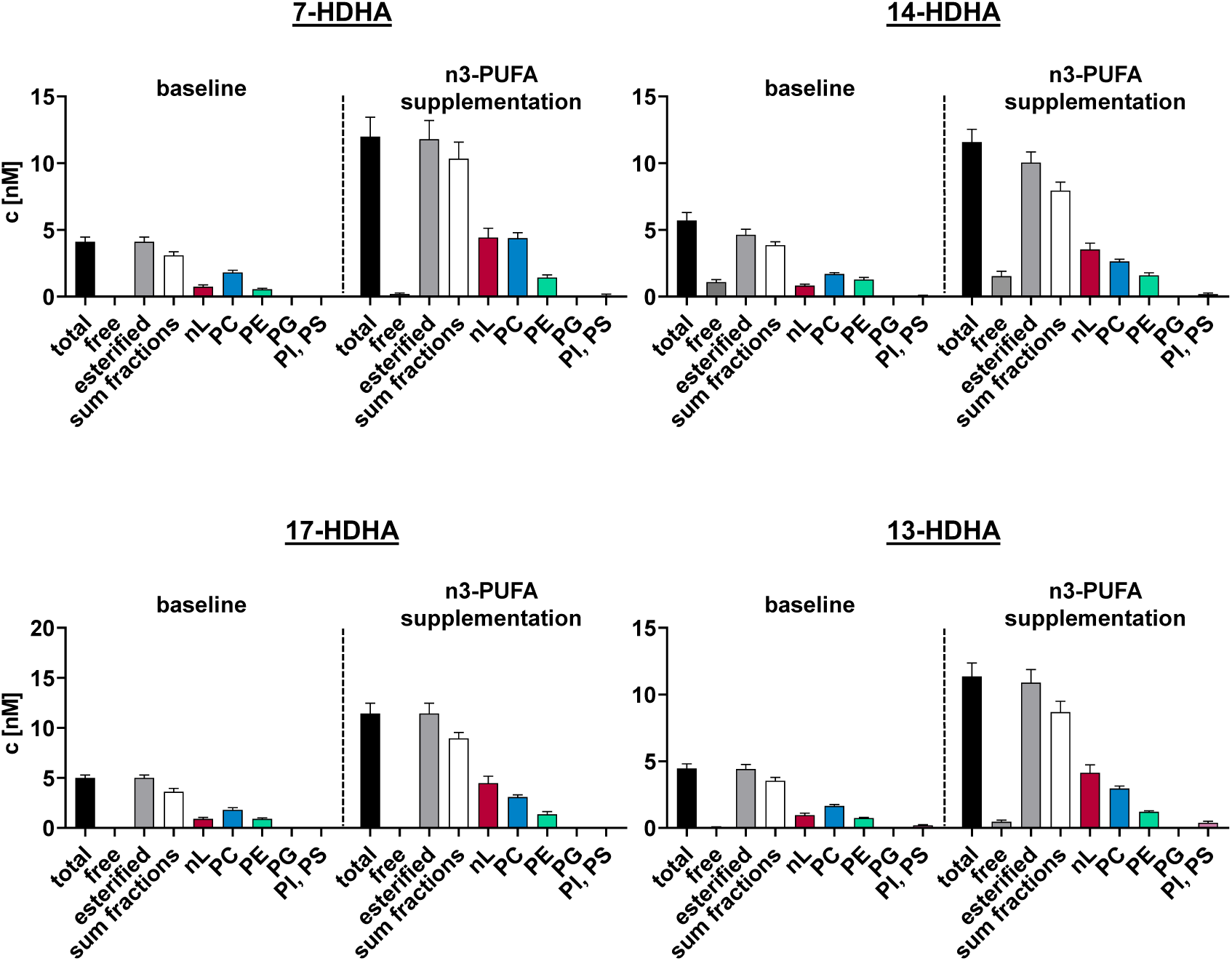
Concentration and distribution of hydroxy-DHA in human plasma esterified in different lipid classes at baseline and after 12 months of n3-PUFA supplementation. Total oxylipins were quantified in the lipid class fractions in plasma of human subjects at baseline and after 12 months of n3-PUFA (1.5 g EPA and 1.8 g DHA/portion; 4 portions per week) supplementation (n = 9). Shown is the concentration ± SE (n = 9) of free and total hydroxy-DHA. All determined concentrations can be found in Table S10. Analysis was carried out using targeted RP-LC-ESI(-)- MS/MS (3–5).

**Fig. S16.**
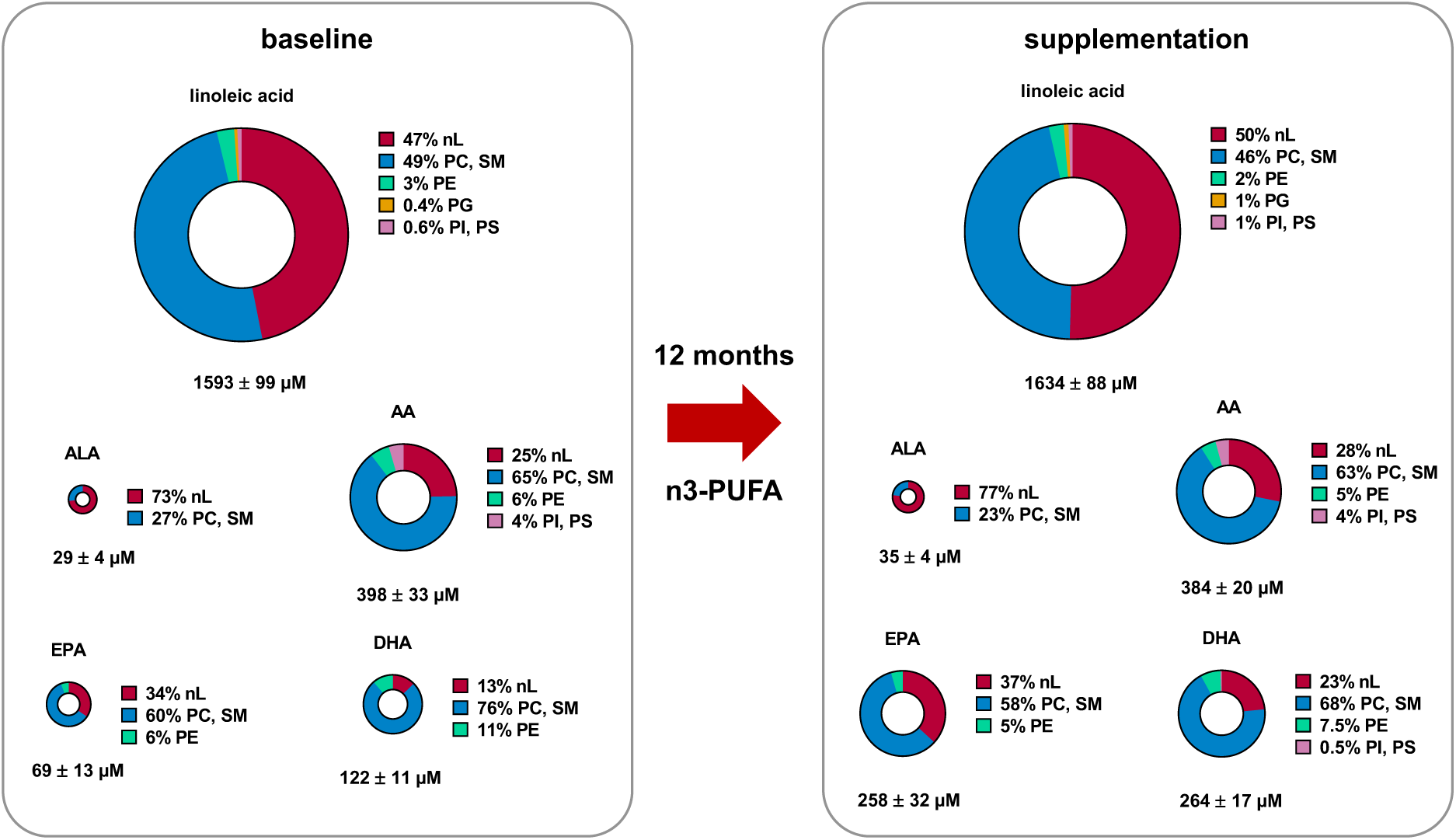
Change of fatty acids esterified in different lipid classes in human plasma following n3-PUFA supplementation. Plasma from 9 human subjects was analyzed at baseline and after 12 months of n3-PUFA (1.5 g EPA and 1.8 g DHA/portion; 4 portions per week) supplementation. Results show concentration and relative lipid class distribution of selected FA in plasma at baseline and after n3-PUFA supplementation. EPA increased from 69 ± 13 µM at baseline to 258 ± 32 µM while the distribution pattern remained comparable. DHA increased from 122 ± 11 µM to 263 ± 17 µM and a shift towards an esterification in nLs was observed. All determined concentrations can be found in Table S10. The areas of the circles reflect the relative concentration of the fatty acids between each other. Analysis was carried out using targeted RP-LC-ESI(-)-MS/MS. Shown are the concentrations ± SE (n = 9) (3–5).

**Fig. S17.**
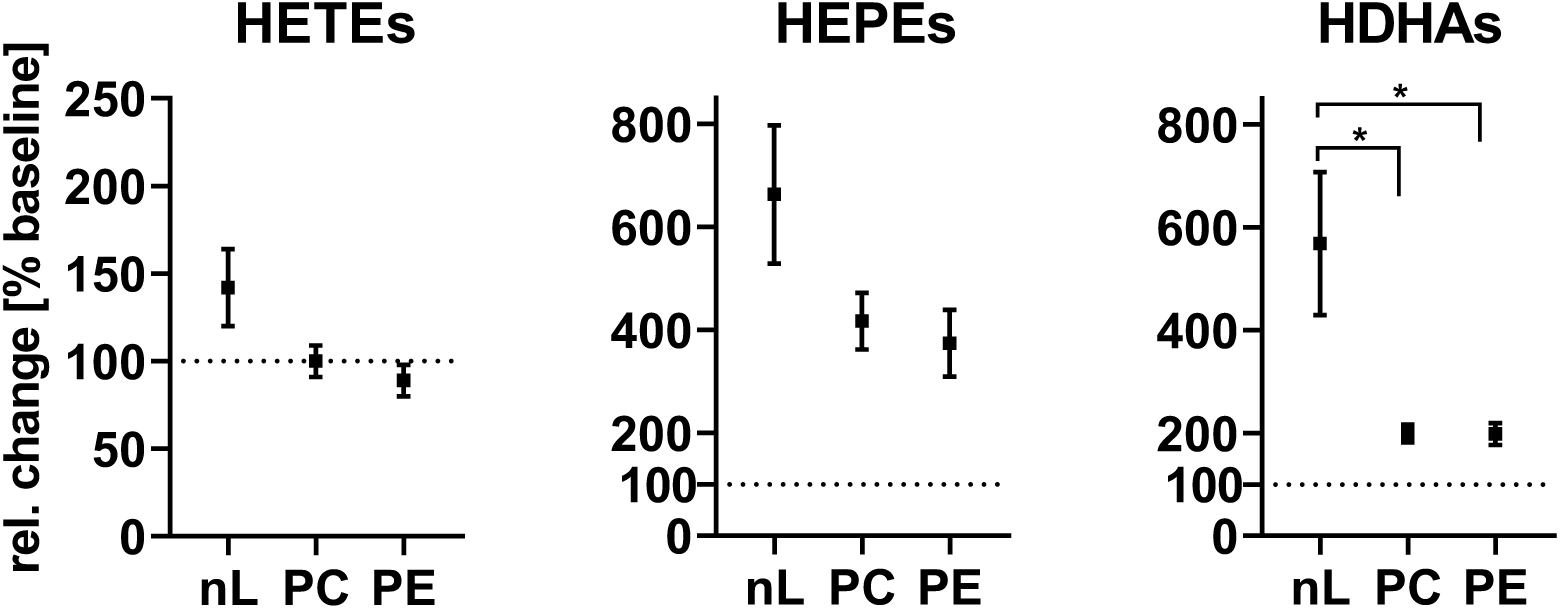
Changes of hydroxy-PUFA concentration in the lipid classes of plasma following 12 months of n3-PUFA supplementation. Total oxylipins were quantified in the lipid class fractions in plasma of human subjects at baseline and after 12 months of n3-PUFA supplementation (1.5 g EPA and 1.8 g DHA per portion; 4 portions per week). Shown are the relative changes in the concentration of the means of all hydroxy-PUFAs of ARA, EPA, and DHA following n3-PUFA supplementation compared to baseline. Differences in the increase in fraction 1 (nLs) vs. the increase in other fractions were elevated by one-way ANOVA followed by Sidak’s multiple comparison test with a statistical significance at p < 0.05 indicated by as *. The concentration of the hydroxy-PUFA and the relative changes can be found in the SI (Tables S10 and S11).

## References

1. Mosblech A, Feussner I, Heilmann I. Oxylipins: structurally diverse metabolites from fatty acid oxidation. Plant Physiol Biochem. 2009;47(6):511–7.

2. Buczynski MW, Dumlao DS, Dennis EA. Thematic Review Series: Proteomics. An integrated omics analysis of eicosanoid biology. J Lipid Res. 2009;50(6):1015–38.

3. Gabbs M, Leng S, Devassy JG, Monirujjaman M, Aukema HM. Advances in Our Understanding of Oxylipins Derived from Dietary PUFAs. Advances in Nutrition. 2015;6(5):513–40.

4. Meirer K, Steinhilber D, Proschak E. Inhibitors of the arachidonic acid cascade: interfering with multiple pathways. Basic Clin Pharmacol Toxicol. 2014;114(1):83–91.

5. Homann J, Lehmann C, Kahnt AS, Steinhilber D, Parnham MJ, Geisslinger G, et al. Chiral chromatography-tandem mass spectrometry applied to the determination of pro-resolving lipid mediators. J Chromatogr A. 2014;1360:150–63.

6. Brash AR. Lipoxygenases: occurrence, functions, catalysis, and acquisition of substrate. J Biol Chem. 1999;274(34):23679–82.

7. Imig JD. Epoxyeicosatrienoic Acids and 20-Hydroxyeicosatetraenoic Acid on Endothelial and Vascular Function. Adv Pharmacol. 2016;77:105–41.

8. Kutzner L, Esselun C, Franke N, Schoenfeld K, Eckert GP, Schebb NH. Effect of dietary EPA and DHA on murine blood and liver fatty acid profile and liver oxylipin pattern depending on high and low dietary n6-PUFA. Food & function. 2020;11(10):9177–91.

9. Gollasch B, Wu G, Dogan I, Rothe M, Gollasch M, Luft FC. Effects of hemodialysis on plasma oxylipins. Physiol Rep. 2020;8(12):e14447.

10. Rund KM, Nolte F, Doricic J, Greite R, Schott S, Lichtinghagen R, et al. Clinical blood sampling for oxylipin analysis - effect of storage and pneumatic tube transport of blood on free and total oxylipin profile in human plasma and serum. Analyst. 2020;145(6):2378–88.

11. Morgan AH, Hammond VJ, Morgan L, Thomas CP, Tallman KA, Garcia-Diaz YR, et al. Quantitative assays for esterified oxylipins generated by immune cells. Nat Protoc. 2010;5(12):1919–31.

12. Rund KM, Peng S, Greite R, Claaßen C, Nolte F, Oger C, et al. Dietary omega-3 PUFA improved tubular function after ischemia induced acute kidney injury in mice but did not attenuate impairment of renal function. Prostaglandins & Other Lipid Mediators. 2020;146:106386.

13. Carpanedo L, Rund KM, Wende LM, Kampschulte N, Schebb NH. LC-HRMS analysis of phospholipids bearing oxylipins. Analytica Chimica Acta. 2024;1326.

14. Schebb NH, Ostermann AI, Yang J, Hammock BD, Hahn A, Schuchardt JP. Comparison of the effects of long-chain omega-3 fatty acid supplementation on plasma levels of free and esterified oxylipins. Prostaglandins Other Lipid Mediat. 2014;113–115:21-9.

15. Annevelink CE, Walker RE, Shearer GC. Esterified Oxylipins: Do They Matter? Metabolites. 2022;12(11).

16. Criscuolo A, Nepachalovich P, Garcia-Del Rio DF, Lange M, Ni Z, Baroni M, et al. Analytical and computational workflow for in-depth analysis of oxidized complex lipids in blood plasma. Nat Commun. 2022;13(1):6547.

17. Hammad LA, Wu G, Saleh MM, Klouckova I, Dobrolecki LE, Hickey RJ, et al. Elevated levels of hydroxylated phosphocholine lipids in the blood serum of breast cancer patients. Rapid Commun Mass Spectrom. 2009;23(6):863–76.

18. Morgan AH, Dioszeghy V, Maskrey BH, Thomas CP, Clark SR, Mathie SA, et al. Phosphatidylethanolamine-esterified eicosanoids in the mouse: tissue localization and inflammation-dependent formation in Th-2 disease. J Biol Chem. 2009;284(32):21185–91.

19. Carpanedo L, Wende LM, Goebel B, Hafner AK, Chromik MA, Kampschulte N, et al. Substrate-dependent incorporation of 15-lipoxygenase products in glycerophospholipids: 15-HETE and 15-HEPE in PI, 17-HDHA in plasmalogen PE, and 13-HODE in PC. J Lipid Res. 2025:100841.

20. Maskrey BH, Bermudez-Fajardo A, Morgan AH, Stewart-Jones E, Dioszeghy V, Taylor GW, et al. Activated platelets and monocytes generate four hydroxyphosphatidylethanolamines via lipoxygenase. J Biol Chem. 2007;282(28):20151–63.

21. Aoyagi R, Ikeda K, Isobe Y, Arita M. Comprehensive analyses of oxidized phospholipids using a measured MS/MS spectra library. J Lipid Res. 2017;58(11):2229–37.

22. Quehenberger O, Dahlberg-Wright S, Jiang J, Armando AM, Dennis EA. Quantitative determination of esterified eicosanoids and related oxygenated metabolites after base hydrolysis. J Lipid Res. 2018;59(12):2436–45.

23. Ostermann AI, Koch E, Rund KM, Kutzner L, Mainka M, Schebb NH. Targeting esterified oxylipins by LC-MS - Effect of sample preparation on oxylipin pattern. Prostaglandins Other Lipid Mediat. 2020;146:106384.

24. Mainka M, Dalle C, Pétéra M, Dalloux-Chioccioli J, Kampschulte N, Ostermann AI, et al. Harmonized procedures lead to comparable quantification of total oxylipins across laboratories. Journal of lipid research. 2020;61(11):1424–36.

25. Fauland A, Trotzmuller M, Eberl A, Afiuni-Zadeh S, Kofeler H, Guo X, et al. An improved SPE method for fractionation and identification of phospholipids. J Sep Sci. 2013;36(4):744–51.

26. Kaluzny MA, Duncan LA, Merritt MV, Epps DE. Rapid separation of lipid classes in high yield and purity using bonded phase columns. J Lipid Res. 1985;26(1):135–40.

27. Pinkart HC, Devereux R, Chapman PJ. Rapid separation of microbial lipids using solid phase extraction columns. J Microbiol Meth. 1998;34(1):9–15.

28. Scholz L, Wende LM, Chromik MA, Kampschulte N, Schebb NH. Oxidative stress leads to the formation of esterified erythro- and threo-dihydroxy-fatty acids in HepG2 cells. Redox Biol. 2025;82:103589.

29. Watanabe S, Souza FDC, Kusumoto I, Shen Q, Nitin N, Lein PJ, et al. Intraperitoneally injected d11-11(12)-epoxyeicosatrienoic acid is rapidly incorporated and esterified within rat plasma and peripheral tissues but not the brain. Prostaglandins Leukot Essent Fatty Acids. 2024;202:102622.

30. Rund KM, Carpanedo L, Lauterbach R, Wermund T, West AL, Wende LM, et al. LC-ESI-HRMS - lipidomics of phospholipids : Characterization of extraction, chromatography and detection parameters. Anal Bioanal Chem. 2024;416(4):925–44.

31. Hartung NM, Mainka M, Pfaff R, Kuhn M, Biernacki S, Zinnert L, et al. Development of a quantitative proteomics approach for cyclooxygenases and lipoxygenases in parallel to quantitative oxylipin analysis allowing the comprehensive investigation of the arachidonic acid cascade. Anal Bioanal Chem. 2023;415(5):913–33.

32. Kutzner L, Rund KM, Ostermann AI, Hartung NM, Galano J-M, Balas L, et al. Development of an Optimized LC-MS Method for the Detection of Specialized Pro-Resolving Mediators in Biological Samples. Frontiers in pharmacology. 2019;10:169.

33. Koch E, Mainka M, Dalle C, Ostermann AI, Rund KM, Kutzner L, et al. Stability of oxylipins during plasma generation and long-term storage. Talanta. 2020;217:121074.

34. Rund KM, Ostermann AI, Kutzner L, Galano JM, Oger C, Vigor C, et al. Development of an LC-ESI(-)-MS/MS method for the simultaneous quantification of 35 isoprostanes and isofurans derived from the major n3- and n6-PUFAs. Anal Chim Acta. 2018;1037:63–74.

35. Koch E, Wiebel M, Hopmann C, Kampschulte N, Schebb NH. Rapid quantification of fatty acids in plant oils and biological samples by LC-MS. Analytical and bioanalytical chemistry. 2021;413(21):5439–51.

36. Browning LM, Walker CG, Mander AP, West AL, Madden J, Gambell JM, et al. Incorporation of eicosapentaenoic and docosahexaenoic acids into lipid pools when given as supplements providing doses equivalent to typical intakes of oily fish. Am J Clin Nutr. 2012;96(4):748–58.

37. Gallego SF, Hermansson M, Liebisch G, Hodson L, Ejsing CS. Total Fatty Acid Analysis of Human Blood Samples in One Minute by High-Resolution Mass Spectrometry. Biomolecules. 2018;9(1).

38. Serafim V, Tiugan DA, Andreescu N, Mihailescu A, Paul C, Velea I, et al. Development and Validation of a LC(-)MS/MS-Based Assay for Quantification of Free and Total Omega 3 and 6 Fatty Acids from Human Plasma. Molecules. 2019;24(2).

39. Moran-Garrido M, Camunas-Alberca SM, Sáiz J, Gradillas A, Taha AY, Barbas C. Deeper insights into the stability of oxylipins in human plasma across multiple freeze-thaw cycles and storage conditions. J Pharmaceut Biomed. 2025;255.

40. Shearer GC, Newman JW. Lipoprotein lipase releases esterified oxylipins from very low-density lipoproteins. Prostaglandins Leukot Essent Fatty Acids. 2008;79(6):215–22.

41. Willenberg I, Ostermann AI, Schebb NH. Targeted metabolomics of the arachidonic acid cascade: current state and challenges of LC–MS analysis of oxylipins. Analytical and Bioanalytical Chemistry. 2015;407(10):2675–83.

42. Sweeley CC. Purification and Partial Characterization of Sphingomyelin from Human Plasma. J Lipid Res. 1963;4:402–6.

43. Quehenberger O, Armando AM, Brown AH, Milne SB, Myers DS, Merrill AH, et al. Lipidomics reveals a remarkable diversity of lipids in human plasma. J Lipid Res. 2010;51(11):3299–305.

44. Ostermann AI, West AL, Schoenfeld K, Browning LM, Walker CG, Jebb SA, et al. Plasma oxylipins respond in a linear dose-response manner with increased intake of EPA and DHA: results from a randomized controlled trial in healthy humans. Am J Clin Nutr. 2019;109(5):1251–63.

45. Schuchardt JP, Ostermann AI, Stork L, Kutzner L, Kohrs H, Greupner T, et al. Effects of docosahexaenoic acid supplementation on PUFA levels in red blood cells and plasma. Prostaglandins, leukotrienes, and essential fatty acids. 2016;115:12–23.

46. Schwalbe-Herrmann M, Willmann J, Leibfritz D. Separation of phospholipid classes by hydrophilic interaction chromatography detected by electrospray ionization mass spectrometry. J Chromatogr A. 2010;1217(32):5179–83.

47. Liebisch G, Vizcaino JA, Kofeler H, Trotzmuller M, Griffiths WJ, Schmitz G, et al. Shorthand notation for lipid structures derived from mass spectrometry. J Lipid Res. 2013;54(6):1523–30.

48. Firl N, Kienberger H, Hauser T, Rychlik M. Determination of the fatty acid profile of neutral lipids, free fatty acids and phospholipids in human plasma. 2013;51(4):799–810.

49. Burdge GC, Wright P, Jones AE, Wootton SA. A method for separation of phosphatidylcholine, triacylglycerol, non-esterified fatty acids and cholesterol esters from plasma by solid-phase extraction. Br J Nutr. 2000;84(5):781–7.

50. Agren JJ, Julkunen A, Penttila I. Rapid separation of serum lipids for fatty acid analysis by a single aminopropyl column. J Lipid Res. 1992;33(12):1871–6.

51. Schebb NH, Kampschulte N, Hagn G, Plitzko K, Meckelmann SW, Ghosh S, et al. Technical recommendations for analyzing oxylipins by liquid chromatography-mass spectrometry. Sci Signal. 2025;18(887):eadw1245.

52. Otoki Y, Metherel AH, Pedersen T, Yang J, Hammock BD, Bazinet RP, et al. Acute Hypercapnia/Ischemia Alters the Esterification of Arachidonic Acid and Docosahexaenoic Acid Epoxide Metabolites in Rat Brain Neutral Lipids. Lipids. 2020;55(1):7–22.

53. Mercola J, D’Adamo CR. Linoleic Acid: A Narrative Review of the Effects of Increased Intake in the Standard American Diet and Associations with Chronic Disease. Nutrients. 2023;15(14).

54. Baker EJ, Miles EA, Burdge GC, Yaqoob P, Calder PC. Metabolism and functional effects of plant-derived omega-3 fatty acids in humans. Progress in Lipid Research. 2016;64:30–56.

55. Kawashima H. Intake of arachidonic acid-containing lipids in adult humans: dietary surveys and clinical trials. Lipids in Health and Disease. 2019;18.

56. Forsyth S, Gautier S, Salem N. Global Estimates of Dietary Intake of Docosahexaenoic Acid and Arachidonic Acid in Developing and Developed Countries. Ann Nutr Metab. 2016;68(4):258–67.

57. Koch E, Löwen A, Schebb NH. Do meals contain a relevant amount of oxylipins? LC-MS-based analysis of oxidized fatty acids in food. Food Chemistry. 2024;438:137941.

58. Koch E, Lowen A, Nikolay S, Willenberg I, Schebb NH. Trans-Hydroxy, Trans-Epoxy, and Erythro-dihydroxy Fatty Acids Increase during Deep-Frying. J Agric Food Chem. 2023;71(19):7508–13.

59. Kato S, Shimizu N, Hanzawa Y, Otoki Y, Ito J, Kimura F, et al. Determination of triacylglycerol oxidation mechanisms in canola oil using liquid chromatography–tandem mass spectrometry. npj Science of Food. 2018;2(1):1.

60. Puca AA, Andrew P, Novelli V, Anselmi CV, Somalvico F, Cirillo NA, et al. Fatty Acid Profile of Erythrocyte Membranes As Possible Biomarker of Longevity. Rejuvenation Research. 2007;11(1):63–72.

61. Raphael W, Sordillo LM. Dietary polyunsaturated fatty acids and inflammation: the role of phospholipid biosynthesis. Int J Mol Sci. 2013;14(10):21167–88.

62. Calder PC. Omega-3 polyunsaturated fatty acids and inflammatory processes: nutrition or pharmacology? Br J Clin Pharmacol. 2013;75(3):645–62.

63. Rise P, Eligini S, Ghezzi S, Colli S, Galli C. Fatty acid composition of plasma, blood cells and whole blood: relevance for the assessment of the fatty acid status in humans. Prostaglandins Leukot Essent Fatty Acids. 2007;76(6):363–9.

64. Haeffner EW, Strosznajder JB. Metabolism of [14C]arachidonic acid-labeled lipids in quiescent and OAG-stimulated ascites tumor cells. Int J Biochem. 1992;24(9):1481–5.

65. Tomita-Yamaguchi M, Babich JF, Baker RC, Santoro TJ. Incorporation, distribution, and turnover of arachidonic acid within membrane phospholipids of B220+ T cells from autoimmune-prone MRL-lpr/lpr mice. J Exp Med. 1990;171(3):787–800.

66. Epand RM. Features of the Phosphatidylinositol Cycle and its Role in Signal Transduction. J Membr Biol. 2017;250(4):353–66.

67. Girton RA, Spector AA, Gordon JA. 15-HETE: Selective incorporation into inositol phospholipids of MDCK cells. Kidney International. 1994;45(4):972–80.

68. Alpert SE, Walenga RW. Human Tracheal Epithelial-Cells Selectively Incorporate 15-Hydroxyeicosatetraenoic Acid into Phosphatidylinositol. Am J Resp Cell Mol. 1993;8(3):273–81.

69. Ramasamy I. Recent advances in physiological lipoprotein metabolism. Clin Chem Lab Med. 2014;52(12):1695–727.

70. Stillwell W, Jenski LJ, Crump FT, Ehringer W. Effect of docosahexaenoic acid on mouse mitochondrial membrane properties. Lipids. 1997;32(5):497–506.

71. de Marco Castro E, Kampschulte N, Murphy CH, Schebb NH, Roche HM. Oxylipin status, before and after LC n-3 PUFA supplementation, has little relationship with skeletal muscle biology in older adults at risk of sarcopenia. Prostaglandins, Leukotrienes and Essential Fatty Acids. 2023;189:102531.

72. Saleh RNM, West AL, Ostermann AI, Schebb NH, Calder PC, Minihane AM. APOE Genotype Modifies the Plasma Oxylipin Response to Omega-3 Polyunsaturated Fatty Acid Supplementation in Healthy Individuals. Frontiers in nutrition. 2021;8:723813.

73. Schmocker C, Zhang IW, Kiesler S, Kassner U, Ostermann AI, Steinhagen-Thiessen E, et al. Effect of Omega-3 Fatty Acid Supplementation on Oxylipins in a Routine Clinical Setting. Int J Mol Sci. 2018;19(1).

74. Schuchardt JP, Ostermann AI, Stork L, Fritzsch S, Kohrs H, Greupner T, et al. Effect of DHA supplementation on oxylipin levels in plasma and immune cell stimulated blood. Prostaglandins, leukotrienes, and essential fatty acids. 2017;121:76–87.

75. Schuchardt JP, Schmidt S, Kressel G, Willenberg I, Hammock BD, Hahn A, et al. Modulation of blood oxylipin levels by long-chain omega-3 fatty acid supplementation in hyper- and normolipidemic men. Prostaglandins, leukotrienes, and essential fatty acids. 2014;90(2-3):27–37.

76. Annika IO, Annette LW, Kirsten S, Lucy MB, Celia GW, Susan AJ, et al. Plasma oxylipins respond in a linear dose-response manner with increased intake of EPA and DHA: results from a randomized controlled trial in healthy humans S1 - 13 M4 - Citavi.

77. Shearer GC, Harris WS, Pedersen TL, Newman JW. Detection of omega-3 oxylipins in human plasma and response to treatment with omega-3 acid ethyl esters. J Lipid Res. 2010;51(8):2074–81.

78. Keenan AH, Pedersen TL, Fillaus K, Larson MK, Shearer GC, Newman JW. Basal omega-3 fatty acid status affects fatty acid and oxylipin responses to high-dose n3-HUFA in healthy volunteers. J Lipid Res. 2012;53(8):1662–9.

79. Koch E, Kampschulte N, Schebb NH. Comprehensive Analysis of Fatty Acid and Oxylipin Patterns in n3-PUFA Supplements. Journal of agricultural and food chemistry. 2022;70(13):3979–88.

80. O’Donnell VB. New appreciation for an old pathway: the Lands Cycle moves into new arenas in health and disease. Biochem Soc Trans. 2022;50(1):1–11.

81. Shearer GC, Newman JW. Impact of circulating esterified eicosanoids and other oxylipins on endothelial function. Curr Atheroscler Rep. 2009;11(6):403–10.

82. Wang L, Gill R, Pedersen TL, Higgins LJ, Newman JW, Rutledge JC. Triglyceride-rich lipoprotein lipolysis releases neutral and oxidized FFAs that induce endothelial cell inflammation. Journal of Lipid Research. 2009;50(2):204–13.

83. Karu N, Kindt A, Lamont L, van Gammeren AJ, Ermens AAM, Harms AC, et al. Plasma Oxylipins and Their Precursors Are Strongly Associated with COVID-19 Severity and with Immune Response Markers. Metabolites. 2022;12(7).

84. Chaves AB, Diniz LS, Santos RS, Lima RS, Oreliana H, Pinto IFD, et al. Plasma oxylipin profiling by high resolution mass spectrometry reveal signatures of inflammation and hypermetabolism in amyotrophic lateral sclerosis. Free Radical Bio Med. 2023;208:285–98.

85. Dalle C, Tournayre J, Mainka M, Basiak-Rasała A, Pétéra M, Lefèvre-Arbogast S, et al. The Plasma Oxylipin Signature Provides a Deep Phenotyping of Metabolic Syndrome Complementary to the Clinical Criteria. International journal of molecular sciences. 2022;23(19).

86. Rund KM, Ostermann AI, Kutzner L, Galano J-M, Oger C, Vigor C, et al. Development of an LC-ESI(-)-MS/MS method for the simultaneous quantification of 35 isoprostanes and isofurans derived from the major n3- and n6-PUFAs. Analytica chimica acta. 2018;1037:63–74.

87. Tsugawa H, Cajka T, Kind T, Ma Y, Higgins B, Ikeda K, et al. MS-DIAL: data-independent MS/MS deconvolution for comprehensive metabolome analysis. Nat Methods. 2015;12(6):523–6.

88. Pi J, Wu X, Feng Y. Fragmentation patterns of five types of phospholipids by ultra-high-performance liquid chromatography electrospray ionization quadrupole time-of-flight tandem mass spectrometry. Analytical Methods. 2016;8(6):1319–32.

## References

1. Browning LM, Walker CG, Mander AP, West AL, Madden J, Gambell JM, et al. Incorporation of eicosapentaenoic and docosahexaenoic acids into lipid pools when given as supplements providing doses equivalent to typical intakes of oily fish. Am J Clin Nutr. 2012;96(4):748–58.

2. Rund KM, Carpanedo L, Lauterbach R, Wermund T, West AL, Wende LM, et al. LC-ESI-HRMS - lipidomics of phospholipids : Characterization of extraction, chromatography and detection parameters. Anal Bioanal Chem. 2024;416(4):925–44.

3. Kutzner L, Rund KM, Ostermann AI, Hartung NM, Galano J-M, Balas L, et al. Development of an Optimized LC-MS Method for the Detection of Specialized Pro-Resolving Mediators in Biological Samples. Frontiers in pharmacology. 2019;10:169.

4. Rund KM, Ostermann AI, Kutzner L, Galano JM, Oger C, Vigor C, et al. Development of an LC-ESI(-)-MS/MS method for the simultaneous quantification of 35 isoprostanes and isofurans derived from the major n3- and n6-PUFAs. Anal Chim Acta. 2018;1037:63–74.

5. Koch E, Mainka M, Dalle C, Ostermann AI, Rund KM, Kutzner L, et al. Stability of oxylipins during plasma generation and long-term storage. Talanta. 2020;217:121074.

6. Carpanedo L, Rund KM, Wende LM, Kampschulte N, Schebb NH. LC-HRMS analysis of phospholipids bearing oxylipins. Analytica Chimica Acta. 2024;1326.

7. Tsugawa H, Cajka T, Kind T, Ma Y, Higgins B, Ikeda K, et al. MS-DIAL: data-independent MS/MS deconvolution for comprehensive metabolome analysis. Nat Methods. 2015;12(6):523–6.

8. Liebisch G, Vizcaino JA, Kofeler H, Trotzmuller M, Griffiths WJ, Schmitz G, et al. Shorthand notation for lipid structures derived from mass spectrometry. J Lipid Res. 2013;54(6):1523–30.

9. Pi J, Wu X, Feng Y. Fragmentation patterns of five types of phospholipids by ultra-high-performance liquid chromatography electrospray ionization quadrupole time-of-flight tandem mass spectrometry. Analytical Methods. 2016;8(6):1319–32.

